# cgDist: An Enhanced Algorithm for Efficient Calculation of pairwise SNP and InDel differences from Core Genome Multilocus Sequence Typing

**DOI:** 10.1101/2025.10.16.682749

**Authors:** Andrea de Ruvo, Pierluigi Castelli, Andrea Bucciacchio, Iolanda Mangone, Verónica Mixão, Vítor Borges, Michele Flammini, Nicolas Radomski, Adriano Di Pasquale

**Affiliations:** Gran Sasso Science Institute, L’Aquila, Italy; Istituto Zooprofilattico Sperimentale dell’Abruzzo e del Molise “G. Caporale”, Teramo, Italy; Instituto Nacional de Saúde Doutor Ricardo Jorge, Lisbon, Portugal

**Keywords:** bacterial genomic surveillance, core genome multilocus sequence typing, outbreak investigation, source attribution, bioinformatics algorithms

## Abstract

Bacterial genomic surveillance requires balancing computational efficiency with genetic resolution for effective outbreak investigation. Traditional cgMLST distance calculations treat all allelic differences as equivalent units, potentially obscuring nucleotide-level variation critical for source attribution. While SNP-based methods provide enhanced resolution, their computational requirements limit routine deployment in surveillance laboratories.

We present cgDist, an algorithm that bridges this resolution gap by calculating nucleotide-level distances directly from cgMLST allelic profiles. The unified cache architecture stores comprehensive alignment statistics, enabling multiple distance calculation modes without redundant computation and supporting both dataset-specific and schema-complete cache generation. This design transforms genomic surveillance from batch processing to continuous streaming analysis, with cumulative performance benefits as laboratories accumulate alignment data.

cgDist functions optimally as a precision “zoom lens” for the detailed investigation of clusters identified through initial cgMLST screening. Rather than restructuring entire population relationships, this targeted approach maximizes epidemiological insight precisely where enhanced resolution is most valuable. The algorithm ensures that cgDist distances are always greater than or equal to corresponding cgMLST distances, preserving epidemiological interpretability while adding critical genetic discrimination.

The system includes integrated recombination detection capabilities that leverage cached alignment statistics to identify potential horizontal gene transfer events through mutation density analysis. This multi-scale analytical framework—from population screening through cluster zoom analysis to recombination detection—provides comprehensive surveillance capabilities within the computational constraints of routine public health practice.

## 1 Introduction

Bacterial genomic epidemiology has been transformed by the advent of next-generation sequencing technologies, enabling high-resolution analysis of pathogen transmission patterns and outbreak investigations [Allard et al., 2018, Didelot et al., 2017, Timme et al., 2017]. This revolution has been particularly impactful for foodborne pathogens such as *Listeria monocytogenes* and *Salmonella enterica*, where accurate source attribution is important for public health protection and outbreak control [Pightling et al., 2018a, Brown et al., 2019].

Core genome multilocus sequence typing (cgMLST) has emerged as the gold standard for bacterial strain typing in surveillance and outbreak investigation workflows [Maiden et al., 2013, Ruppitsch et al., 2015a, European Food Safety Authority, 2022]. This approach characterizes bacterial isolates using allelic profiles derived from conserved core genome loci, providing a standardized framework that balances discriminatory power with computational tractability [Silva et al., 2018, Moura et al., 2016]. The method’s success stems from its ability to generate comparable results across different laboratories while maintaining sufficient resolution for epidemiological applications [Moura et al., 2016, Ruppitsch et al., 2015b].

Traditional cgMLST analysis relies on allelic difference calculations, for instance using Hamming distance, which counts the number of differing alleles between isolate pairs [Seemann, 2020]. While computationally efficient and widely adopted, this approach treats all allelic differences as equivalent units, regardless of the underlying nucleotide-level variation, potentially limiting the resolution for accurate outbreak reconstruction [Henri et al., 2017, Ragon et al., 2008, Morganti et al., 2018].

Single nucleotide polymorphism (SNP)-based methods provide the highest phylogenetic resolution by examining individual nucleotide differences within genomic regions shared across all isolates (i.e., the core genome) [Schürch et al., 2018, Davis et al., 2015, Gardner et al., 2015]. These approaches process raw sequencing data or assembled genomes to construct SNP matrices that accurately reflect evolutionary relationships and transmission patterns. However, such methods require computational resources and complex bioinformatics pipelines, limiting their applicability for routine surveillance where rapid turnaround times are required [Deneke et al., 2021, Uelze et al., 2020, Mixão et al., 2025].

The computational challenges facing genomic surveillance systems are further compounded by Amdahl’s law, which demonstrates that excessive parallelization cannot overcome fundamental algorithmic bottlenecks due to thread synchronization overhead and memory access coordination [Amdahl, 1967]. This limitation is particularly relevant for real-time surveillance applications, where processing thousands of isolates with rapid turnaround requirements demands both algorithmic efficiency and scalable performance characteristics [Uelze et al., 2020, Timme et al., 2017].

Current genomic surveillance infrastructure faces a gap between the computational efficiency required for routine monitoring and the resolution necessary for accurate epidemiological inference. An ideal solution would leverage precomputed allelic profiles to minimize computational overhead while incorporating nucleotide-level information to enhance discriminatory power, remaining independent of the specific allele calling methodology to ensure compatibility across different surveillance platforms and laboratory workflows. Such an approach would enable rapid integration of new isolates without genome reprocessing, ensuring timely detection of emerging transmission clusters and facilitating effective public health interventions [Allard et al., 2018].

An additional complexity in outbreak cluster delineation arises from horizontal gene transfer and genetic recombination events, which can inflate genetic distances between epidemiologically related isolates and disrupt cluster boundaries [Arnold et al., 2018, Holden et al., 2004, Feil et al., 2003]. These genetic exchanges can cause closely related isolates to appear more distant than expected, potentially fragmenting outbreak clusters or merging unrelated transmission chains [Didelot and Maiden, 2010, Croucher et al., 2015]. Integration of recombination detection into clustering workflows could improve the accuracy of outbreak delineation by identifying loci affected by horizontal transfer, enabling more robust cluster assignments that reflect true epidemiological relationships rather than confounding evolutionary signals.

Here, we present cgDist, an algorithm that combines the computational efficiency of cgMLST with the enhanced resolution of SNP-based methods for bacterial genomic surveillance. Additionally, we demonstrate how cgDist’s unified cache architecture naturally enables integrated recombination analysis, providing a comprehensive multi-scale approach from population-level screening to detailed genetic characterization.

## 2 Methods

**Terminology clarification:** The term “distances” is used in a computational context in this manuscript and refers to genomic differences between pairs of genomes, rather than phylogenetic distances that represent evolutionary time. Additionally, “cgMLST distance” refers to the traditional allelic Hamming distance calculation used in cgMLST analysis, where each allelic difference contributes one unit regardless of the underlying sequence variation.

### 2.1 Algorithm Design and Implementation

#### 2.1.1 cgDist Algorithm: Conceptual Framework

**Basic Idea** The fundamental concept behind cgDist is straightforward: instead of treating all genetic differences as equal units, we examine the actual DNA sequences using cgMLST data to count the precise number of mutations between bacterial samples. Traditional cgMLST analysis assigns each gene variant a unique identifier (like a barcode) and calculates distances by simply counting how many identifiers differ between samples— this is called allelic Hamming distance, which we use throughout this manuscript to refer to traditional cgMLST distance calculations. While this approach provides standardized and reproducible results, it cannot distinguish between alleles with minimal sequence variation and those with substantial nucleotide differences.

In contrast, when two samples have different barcodes for the same gene, cgDist retrieves the actual DNA sequences behind those barcodes and aligns them to count the exact number of single nucleotide polymorphisms (SNPs) and insertion-deletion events (InDels). For example, if traditional analysis shows “2 allele differences” between two bacterial isolates, cgDist might reveal this actually represents “5 SNPs and 2 InDel events”— providing much finer resolution for distinguishing closely related outbreak strains from unrelated background isolates (Figure 1). This enhanced precision is particularly valuable in food safety investigations where accurate source attribution can prevent widespread contamination [Allard et al., 2018, Pightling et al., 2018a], and in hospital settings where distinguishing true transmission chains from coincidental similarities is crucial for infection control [Moura et al., 2017, Gymoese et al., 2017].

**Figure 1.**
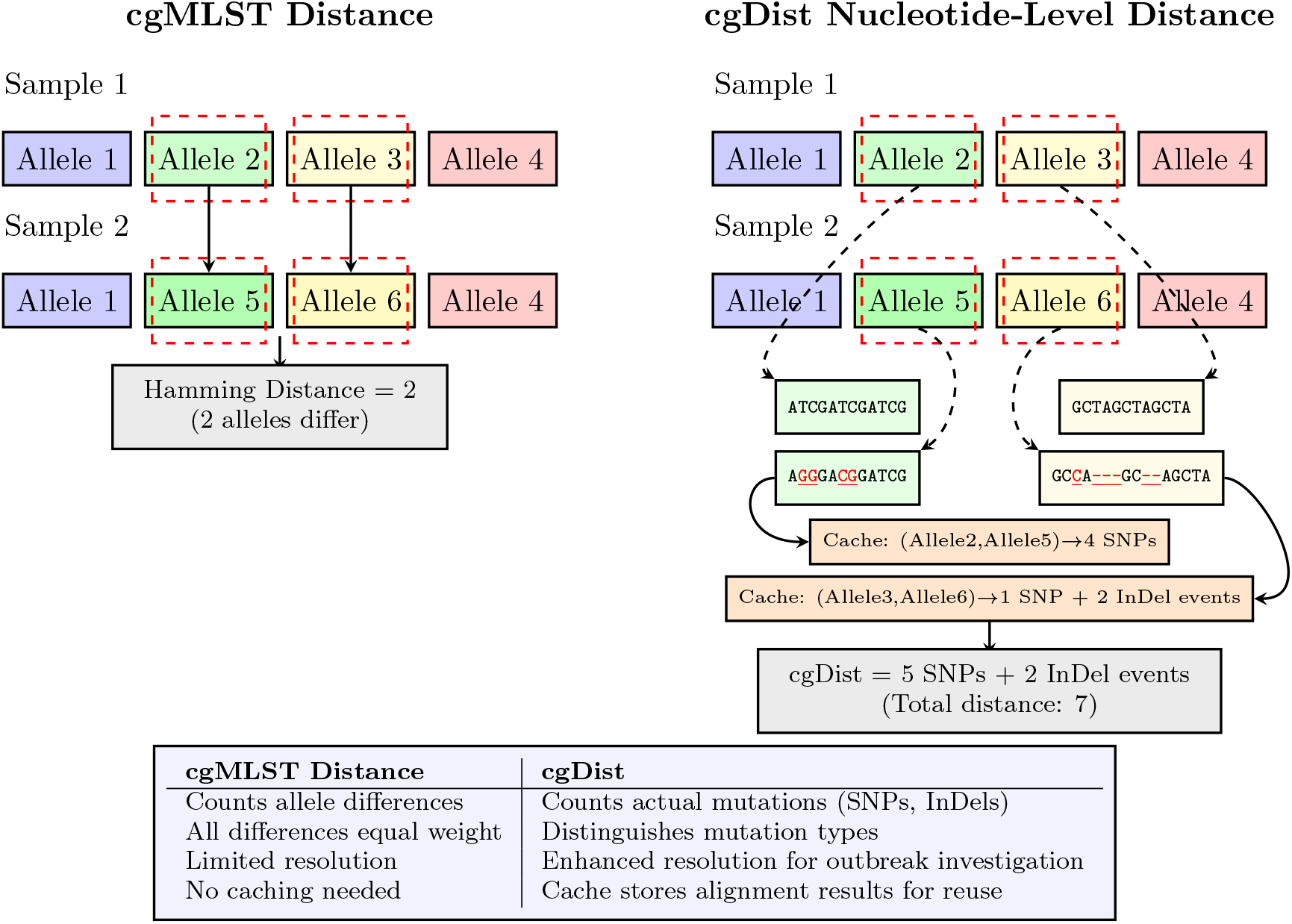
Comparison of distance calculation approaches in bacterial genomics. The figure shows two methods: cgMLST distance (left) counts genes with different allele identifiers, treating all differences equally (distance = 2 in this example). cgDist (right) examines the actual DNA sequences behind differing alleles to quantify both SNPs and InDel events, revealing that two allele differences actually represent 5 SNPs and 2 InDel events (total distance = 7). The cache system stores alignment results (orange boxes) for performance optimization, demonstrating how cgDist intuitively provides higher resolution than cgMLST distance (7 vs. 2) for epidemiological investigations.

#### 2.1.2 Algorithm Implementation

cgDist is implemented in Rust, leveraging the language’s guarantees of memory safety [Matsakis and Klock, 2014] and high-performance execution model, to compute nucleotide-level distances directly from cgMLST allelic profiles through sequence-level analysis. Unlike traditional allele-based distance calculations that treat all allelic differences as equivalent units [Seemann, 2020], cgDist performs pairwise sequence alignment using the Needleman-Wunsch global alignment algorithm [Needleman and Wunsch, 1970] via the Parasail library (parasail-rs v0.3.5 Rust bindings) [Daily, 2016] to quantify actual SNP and insertion-deletion (InDel) differences between alleles. This approach provides enhanced genomic clustering resolution for surveillance applications while maintaining the computational advantages of working with cgMLST data, allowing incremental analysis without reprocessing entire datasets from raw sequencing data [Silva et al., 2018]. Technical implementation details and debugging capabilities are described in Supplementary Methods Sections 7 and 7.

**Distance Calculation Modes** cgDist supports four complementary distance calculation modes, each designed for specific epidemiological applications. The **SNPs-only** mode counts single nucleotide polymorphisms exclusively, providing the finest resolution for highly similar isolates. The **SNPs + InDel events** mode combines SNPs with the number of distinct insertion-deletion events, where each continuous gap region is counted as a single mutational event regardless of length. The **SNPs + InDel bases** mode counts SNPs plus the total number of inserted or deleted nucleotides, providing comprehensive nucleotide-level divergence measurement. The **Hamming** mode computes traditional cgMLST allelic differences for baseline comparison and validation.

**Unified Distance Cache Architecture and Memoization** cgDist implements a unified distance cache system that stores all alignment statistics (SNPs, InDel events, InDel bases) in a single cache entry, enabling alignment reuse across different analysis configurations for the same dataset. The system implements intelligent memoization by scanning allelic profiles to identify only unique allele pairs requiring alignment computation, eliminating redundant calculations through the symmetric property of pairwise distances. This design exploits the characteristic clonal population structures of bacterial surveillance datasets [Moura et al., 2016], where many samples share identical alleles at multiple loci, to minimize computational redundancy.

This architecture provides performance benefits: (1) switching between distance calculation modes without recalculation, (2) varying sample completeness thresholds (*θ*_*s*_) and loci completeness thresholds (*θ*_*ℓ*_) requires only matrix filtering without recomputing alignments, (3) incremental processing enables efficient integration of new isolates without reprocessing historical data, and (4) cumulative cache benefits within species enable progressive performance gains as surveillance programs accumulate alignment data over time.

The cache employs a two-level architecture with hash-based keys for rapid lookup and compressed binary storage for efficient disk usage. Each cache entry maps a locus-allele pair tuple to comprehensive alignment statistics including SNP count, InDel events, InDel bases, and computation timestamp. Cache metadata tracks alignment parameters, hasher type, and compatibility information to prevent misuse across different configurations [Deneke et al., 2021].

The cache architecture supports two operational modes: dataset-specific caching where only allele pairs present in the current analysis are computed and stored, and schema-complete caching where all possible allele pair combinations within a cgMLST schema are pre-computed independently of any specific dataset. While schema-complete mode would enable processing all allele sequences from schema FASTA files to generate exhaustive alignment statistics for all possible allele pairs, this comprehensive pre-computation approach requires substantial computational resources and storage capacity, and is superfluous for the objectives of the current study. Therefore, it is not explored here but reserved for future investigation.

**Input Data Requirements and Output Formats** cgDist requires three primary inputs: (1) a cgMLST allelic profiles matrix representing the sample dataset, where rows correspond to genomic samples and columns to cgMLST loci, with each cell containing the hash identifier for the corresponding allele, (2) a cgMLST schema consisting of FASTA files containing nucleotide sequences for all alleles, and (3) the distance calculation mode specification. An optional cache file containing pre-computed alignment statistics can be provided to accelerate computation by reusing previously calculated distances.

The allelic profiles matrix serves as the computational roadmap and defines the sample dataset for analysis, with hash identifiers acting as keys to retrieve sequences from the schema. cgDist supports multiple hashing algorithms with a flexible architecture for custom hashers, requiring only that hash values establish bijective mapping between matrix entries and schema sequences.

cgDist generates multiple output formats: (1) symmetric distance matrices in standard formats (TSV, CSV, PHYLIP, NEXUS), (2) updated cache files containing newly computed alignment statistics for future reuse, (3) detailed computation logs with performance metrics and cache utilization statistics, (4) cache files with embedded metadata documenting analysis parameters and data provenance, and (5) optional human-readable alignment details enabling researchers to manually inspect specific nucleotide differences contributing to distance calculations, validate alignment quality, and identify potential sequencing or assembly artifacts. All outputs maintain sample identifier consistency with the input matrix and support integration into downstream phylogenetic analysis workflows.

**Core Algorithm and Execution Flow** The cgDist computation follows a three-phase approach designed for optimal performance and scalability (Algorithm 1, Figure 2). Phase 1 performs profile filtering based on user-defined completeness thresholds for samples (*θ*_*s*_: minimum fraction of loci required per sample) and loci (*θ*_*ℓ*_: minimum fraction of samples required per locus), enabling rapid exploration of different analytical stringency levels without recomputing alignments. To ensure epidemiologically meaningful results, the algorithm requires that sample pairs share a minimum number of loci (min loci) before distances are computed, preventing unreliable estimates when too few loci are in common [Didelot and Falush, 2007, Ruppitsch et al., 2015b]. Phase 2 implements cache-aware precomputation, extracting unique allele pairs and performing batch alignment only for missing cache entries. Phase 3 assembles the distance matrix using cached alignment statistics with parallel execution across sample pairs.

**Figure 2.**
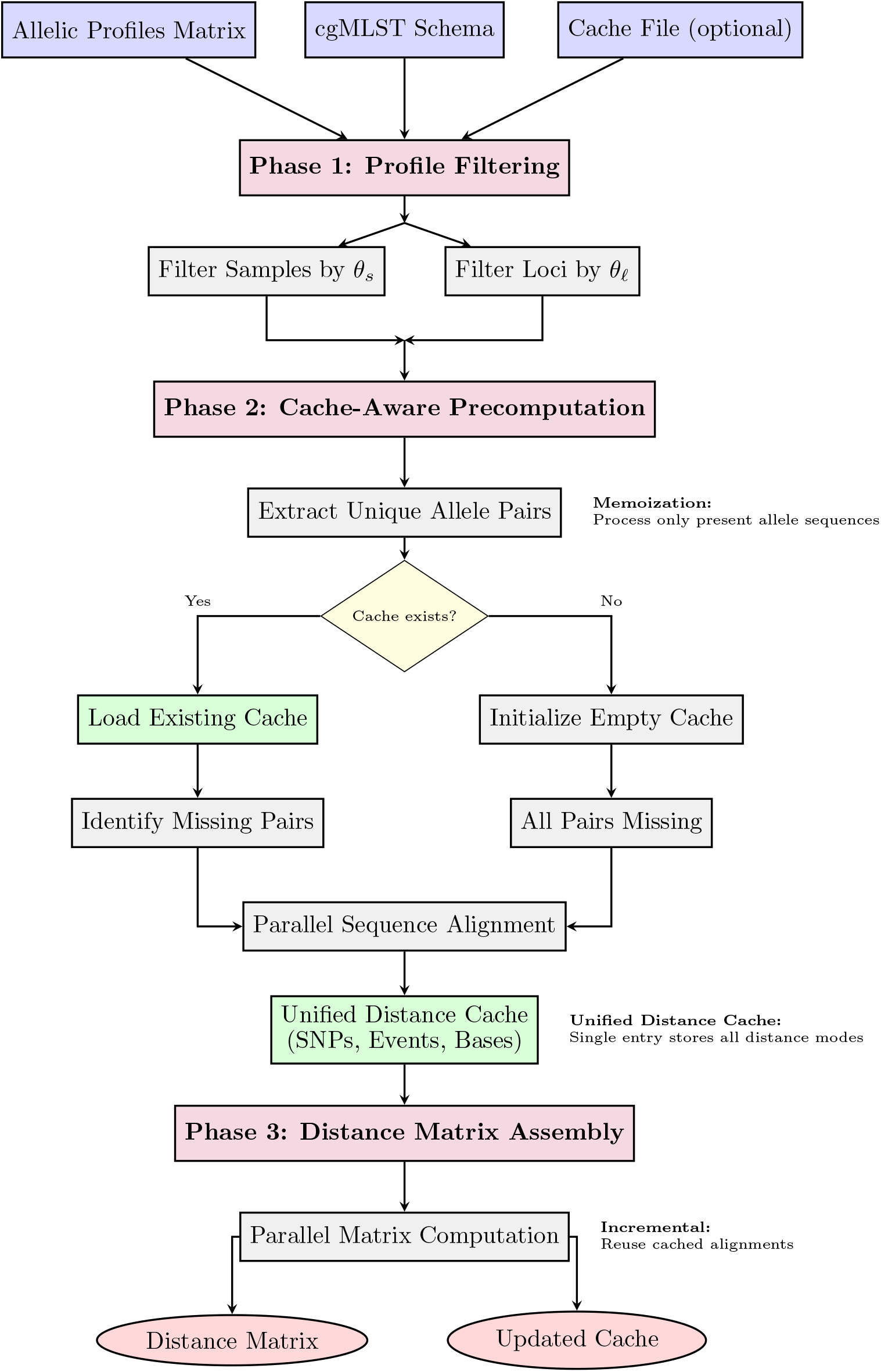
cgDist algorithm workflow showing the three-phase approach with cache-aware optimization. Phase 1 filters input profiles based on quality thresholds (*θ*_*s*_ for sample completeness, *θ*_*ℓ*_ for locus completeness). Phase 2 implements memoization by extracting only unique allele pairs present in the dataset and leverages existing cache when available. Phase 3 assembles the distance matrix using parallel computation with cached alignment statistics. The unified distance cache architecture stores all distance calculation modes (SNPs, InDel events, InDel bases) in single entries, enabling rapid mode switching and incremental processing across surveillance runs.

A critical design invariant ensures that cgDist values are always greater than or equal to corresponding cgMLST distances (*d*_cgMLST_ ≤ *d*_cgDist_). In SNPs-only mode, when sequence alignment reveals zero SNPs for alleles with different hash identifiers (indicating only InDel differences), the algorithm applies a Hamming fallback mechanism, contributing +1 to maintain the ordering constraint while preserving epidemiological consistency.

##### Algorithm 1: cgDist Core Algorithm with Unified Distance Cache

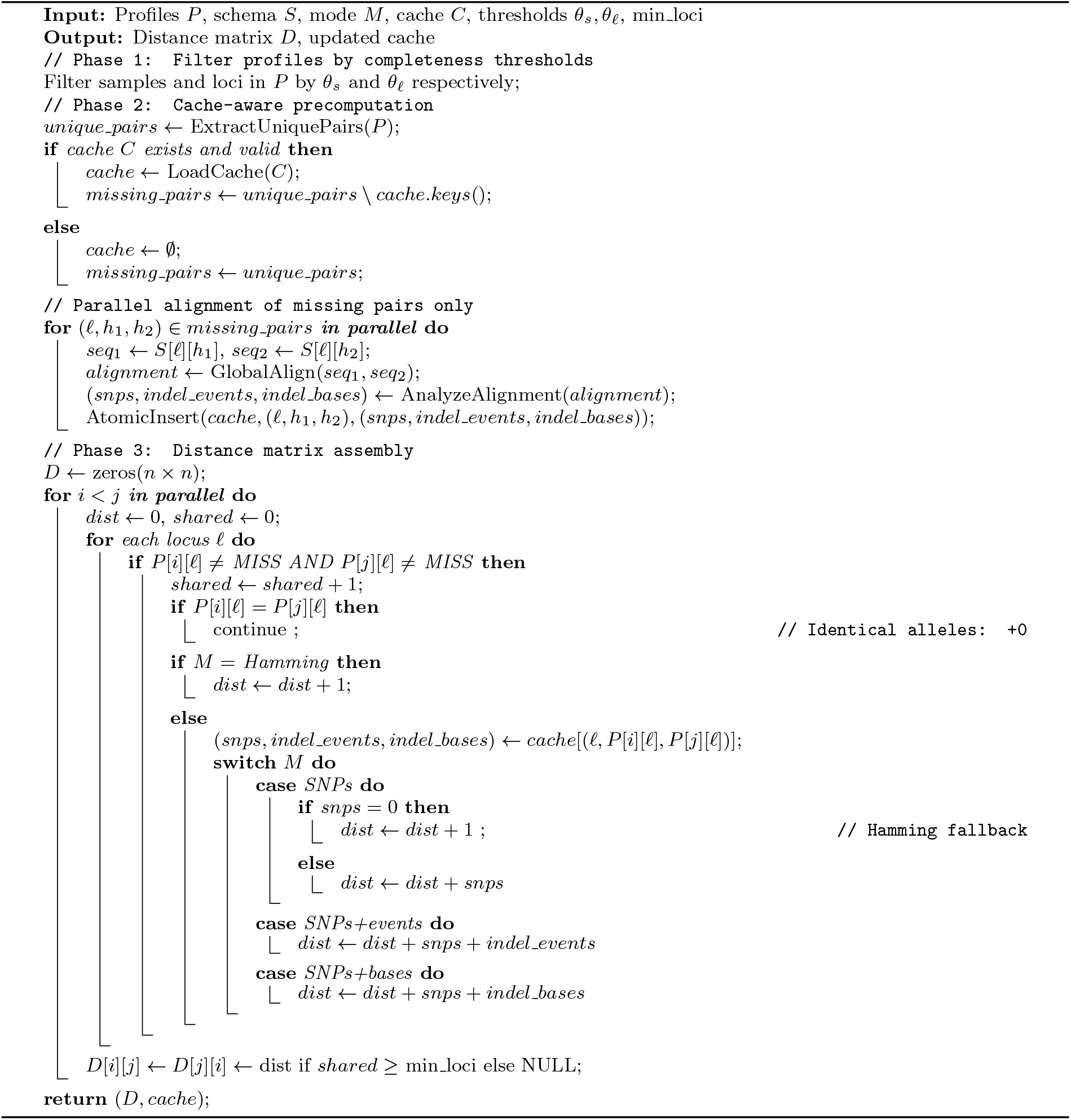

**Distance Calculation Algorithm** The core distance calculation proceeds through sequence alignment analysis to extract nucleotide-level differences between allele pairs. For each unique allele pair identified in the profiles matrix, cgDist retrieves the corresponding nucleotide sequences from the schema and performs global sequence alignment using the Parasail library. The alignment results are then analyzed character-by-character to identify and classify genetic variants into three categories: single nucleotide polymorphisms (SNPs), insertion-deletion events (discrete gap regions), and insertion-deletion bases (total nucleotides involved in gaps).

The algorithm maintains running counters for each variant type while processing the alignment, ensuring consistent classification of complex genomic variations. Gap tracking logic distinguishes between continuous indel regions and separate events, enabling accurate event counting that reflects the underlying mutational processes. Different distance calculation modes selectively combine these counters to generate the final distance metrics according to the specified analysis requirements.

The detailed algorithm for analyzing pairwise sequence alignments and computing different distance metrics is provided in Supplementary Methods Section 7.

**Alignment Parameters and Mode Selection** For this study, we employed cgDist’s DNA-strict alignment mode to ensure conservative, high-confidence distance measurements. This approach builds upon established global alignment principles [Needleman and Wunsch, 1970] with parameters optimized for bacterial genomic sequences (detailed parameters in Supplementary Methods Section 7).

#### 2.1.3 Mathematical Foundation: cgMLST-cgDist Ordering Theorem

We establish the fundamental mathematical relationship between cgMLST distance calculations and nucleotide-level distance calculations through formal analysis.

##### Theorem 1

(cgMLST-cgDist Ordering). *For any two genomic samples S*_*i*_ *and S*_*j*_ *analyzed using the same cgMLST scheme:*

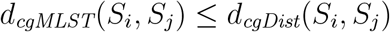

To prove this theorem, we first establish a supporting lemma:

##### Lemma 1.

*For any locus ℓ where alleles differ* 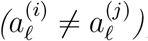, *the number of nucleotide-level mutations satisfies* 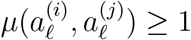.

*Proof of Lemma 1*. Since alleles 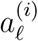 and 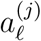 have different identifiers, their underlying DNA sequences must differ by at least one nucleotide position. This difference manifests as either: (1) one or more single nucleotide polymorphisms (SNPs), (2) one or more insertion-deletion events (InDels), or (3) a combination of both. In all cases, 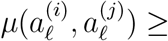.

For the special case of SNPs-only mode where sequence alignment reveals zero SNPs for alleles with different identifiers (indicating InDel-only variation), cgDist applies Hamming fallback: 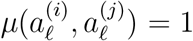. This design choice preserves the fundamental inequality when SNP analysis cannot capture underlying sequence variation.

*Proof of Theorem 1*. Let 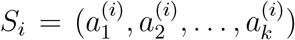 and 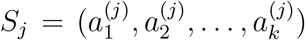 be allelic profiles across *k* cgMLST loci. Define 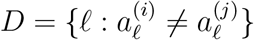 as the set of differing loci.

By definition, the cgMLST distance is: *d*_cgMLST_(*S*_*i*_, *S*_*j*_) = |*D*|.

The cgDist distance counts nucleotide-level mutations: 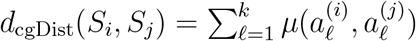, where 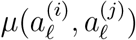 represents the number of mutations between alleles at locus *ℓ*.

Note that for loci where alleles are identical (*ℓ* ∉ *D*), we have 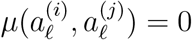.

Therefore:

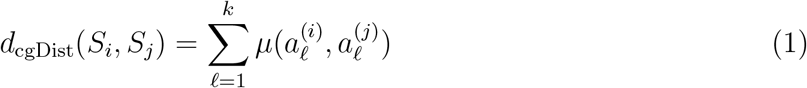

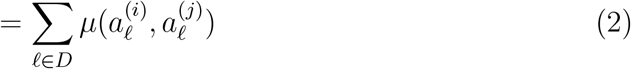

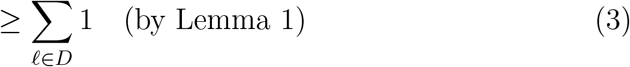

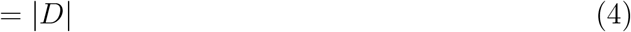

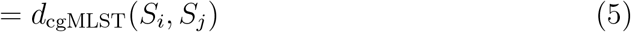

This theorem guarantees that cgDist provides a conservative refinement of traditional cgMLST distance calculation under all computational scenarios, never underestimating evolutionary distance while revealing hidden mutational complexity when sequence alignment succeeds.

##### Corollary 1.

*In SNPs-only mode, equality d*_*cgMLST*_(*S*_*i*_, *S*_*j*_) = *d*_*cgDist*_(*S*_*i*_, *S*_*j*_) *holds when each differing allele pair either: (1) contains exactly one SNP (regardless of InDel presence), or (2) contains zero SNPs, triggering the Hamming fallback mechanism*. □

**Computational Complexity and Performance** cgDist exhibits time complexity *O*(*n*^2^ *× s*^2^ *× l*) for computing pairwise distances between *n* samples across *l* loci with maximum sequence length *s*. Cache optimization reduces this to *O*(*n*^2^ *× l*) when alignment results are reused. Space complexity is *O*(*n*^2^ + *l × u*^2^) where *u* represents average unique alleles per locus, with the dominant term depending on dataset characteristics. Detailed complexity analysis including bounds, optimizations, and practical performance considerations is provided in Supplementary Methods Section 7.

**Deterministic Execution and Reproducibility** cgDist guarantees deterministic results through hash-based allele identification, canonical cache keys using ordered hash pairs, deterministic sequence alignment, and parallel execution safety via read-only cache operations during matrix computation. Given identical inputs, cgDist produces identical distance matrices regardless of execution environment, thread count, or processing order, ensuring scientific reproducibility and regulatory compliance [Allard et al., 2018, Timme et al., 2017].

**Extensibility and Schema Flexibility** cgDist implements a flexible architecture that enables adaptation to diverse cgMLST schemas and laboratory workflows. The system supports multiple hashing algorithms with a plugin system for implementing custom hashers to accommodate specialized applications or legacy compatibility requirements.

The flexible architecture enables laboratories to implement domain-specific hashing strategies, maintain compatibility with existing tools, or meet specific regulatory requirements while leveraging cgDist’s performance optimizations. Custom hashers can be designed for specialized organisms, experimental research applications, or integration with institutional bioinformatics pipelines.

Additionally, cgDist provides both command-line interfaces and programmatic APIs for integration into automated workflows, supporting multiple input/output formats and configuration management. This flexibility ensures compatibility with existing laboratory information management systems and enables seamless integration into diverse surveillance infrastructure without disrupting established analytical workflows.

### 2.2 Experimental Design and Validation

#### 2.2.1 Datasets and Experimental Setup

**Validation Datasets** We validated cgDist using four comprehensive datasets from the OHEJP BeONE project [Mixão et al., 2025]: 1,426 and 1,874 *Listeria monocytogenes* isolates (referred to as Lm-1426 and Lm-1874 datasets respectively) and 1,434 and 1,540 *Salmonella enterica* isolates (referred to as Se-1434 and Se-1540 datasets respectively) from European surveillance networks spanning multiple years and geographic regions. We used *Listeria monocytogenes* and *Salmonella enterica* as representative clonal and panmictic models, respectively, reflecting the low recombination rates observed in *Listeria* [Orsi et al., 2008, Cai et al., 2008] and the high recombination activity reported in *Salmonella* [Diez-Villasenor and Rodriguez-Valera, 2021, *Gonzalez-Escalona et al*., *2021]. All four datasets were used for comprehensive validation, mathematical constraint verification, and performance benchmarking*.

*The selection of these bacterial species and dataset pairs provides complementary validation perspectives across different population genetic structures and sampling strategies. L. monocytogenes* exhibits predominantly clonal population structure with limited recombination, while *S. enterica* displays panmictic characteristics with more frequent horizontal gene transfer, enabling assessment of cgDist performance across the spectrum of bacterial population genetics. For each species, two datasets with distinct sampling designs were used: (i) datasets with comprehensive source attribution metadata (Lm-1426, Se-1540) representing sampling diversity across BeONE consortium partners, where isolate selection was determined by partner contributions and outbreak investigation priorities; and (ii) datasets without detailed source metadata (Lm-1874, Se-1434) representing species diversity from international public archives (ENA/SRA), providing broader coverage of genetic variation present in surveillance databases. This dual-dataset approach per species enables validation across both targeted surveillance collections and broader genomic diversity landscapes.

For epidemiological source prioritization analysis, we utilized the two datasets with comprehensive source attribution metadata: the 1,426 *L. monocytogenes* dataset comprising isolates from clinical surveillance (76.4%), processing plant monitoring (14.1%), food samples (6.9%), and other sources, spanning 28 distinct sequence types with ST5 and ST6 predominant; and the 1,540 *S. enterica* dataset including isolates from clinical (59.0%), environmental (3.1%), animal (5.8%), and other sources (32.1%), representing more than 75 sequence types with ST11, ST34, and ST19 most prevalent. The remaining two datasets (1,874 *Listeria* and 1,434 *Salmonella*) lack detailed source classification metadata and were therefore used exclusively for validation and performance testing.

Allelic profiles were generated using chewBBACA v3.3.10 [Silva et al., 2018] with the `--hash-profiles crc32` parameter. We used the INNUENDO cgMLST schemas [Moura et al., 2017], available through ChewieNS [Machado et al., 2020] for *L. monocytogenes* (accessed March 2024) and *S. enterica* (accessed March 2024). For *S. enterica* analysis, we applied the EFSA-filtered loci subset (3,255 loci) following European food safety surveillance standards [European Food Safety Authority, 2022] to ensure compatibility with regulatory frameworks and established epidemiological thresholds. All assemblies met quality thresholds for reliable cgMLST analysis, and distance calculations satisfied the fundamental constraint *d*_cgMLST_ ≤ *d*_cgDist_ across all datasets.

Mathematical validation, correlation analysis, and performance benchmarking were conducted using all available datasets. For epidemiological analyses including source prioritization and clustering resolution assessment, we applied a 98% completeness threshold (*θ*_*s*_ = 0.98) requiring samples to have allele calls for at least 98% of the cgMLST loci. This threshold is consistent with established cgMLST quality control practices for *L. monocytogenes*, where high-quality sequences typically achieve completeness levels above 98.5% [Palma et al., 2022]. For *S. enterica*, we applied the same stringent threshold, though this represents a more conservative filtering approach than typically required for *Salmonella* surveillance. This quality filtering is required for epidemiological analysis where incomplete allelic profiles could introduce noise in outbreak investigation decisions.

**Comparative Analysis Framework** For both species, we performed parallel distance calculations using cgmlst-dists for cgMLST distances [Seemann, 2020] and cgDist in both SNPs-only and SNPs+InDel-events modes. Both cgDist modes were selected to provide complementary perspectives on epidemiological source prioritization analysis: SNPs-only mode provides a focused analysis of nucleotide substitutions, while SNPs+InDel-events mode provides an optimal balance between resolution enhancement and epidemiological interpretability by capturing discrete mutational events without being influenced by the length heterogeneity that characterizes SNPs+InDel-bases mode. This framework enables direct assessment of resolution improvement and epidemiological utility across identical sample sets, with particular focus on epidemiologically relevant distance ranges (≤7 alleles for *L. monocytogenes* [Moura et al., 2016], ≤ 10 alleles for *S. enterica* [European Food Safety Authority, 2022, Radomski et al., 2019]).

#### 2.2.2 Validation and Analysis Methods

**Mathematical Validation and Statistical Analysis** We implemented systematic, schema-independent validation of the core design constraint *d*_cgMLST_ ≤ *d*_cgDist_ by computing both distance types for all pairwise comparisons across diverse cgMLST schemas and bacterial species. This validation framework is inherently schema-independent, requiring no prior knowledge of locus selection, allelic diversity, or species-specific genetic architecture. Any constraint violations are flagged for detailed investigation, including examination of underlying allelic profiles and sequence alignments. The validation framework also confirms correct implementation of the Hamming fallback mechanism for cases where sequence alignment reveals zero SNPs for alleles with different hash identifiers.

The schema-independent nature of our validation approach ensures that the mathematical guarantees established for cgDist apply universally across any cgMLST scheme, regardless of the specific loci included, target organism, or laboratory implementation. This universality is achieved through the fundamental property that the ordering relationship is preserved at the sequence level for any pair of alleles, independent of their biological context or schema design principles.

**Statistical Correlation Analysis** To assess whether cgDist maintains consistent relationships with cgMLST distances while providing enhanced resolution, we computed both Pearson correlation coefficients [Pearson, 1895] and Spearman rank correlation [Spearman, 1904] between distance methods. Pearson correlation measures linear relationships between the actual distance values, evaluating whether cgDist distances scale proportionally with cgMLST distances. Spearman rank correlation assesses the preservation of relative ordering between isolate pairs, which is critical for epidemiological clustering where rank-based relationships determine outbreak linkage decisions.

**Multi-scale Clustering Analysis Framework** To evaluate cgDist’s utility across different epidemiological scales—from global population-level analysis to detailed outbreak investigation—we implemented a comprehensive three-tiered clustering assessment framework. This multi-scale approach validates cgDist’s resolution enhancement capabilities: (1) **Population screening**—initial cgMLST analysis to identify epidemiologically relevant clusters and potential outbreak groups; (2) **Cluster zoom analysis**—cgDist nucleotide-level distance calculation on selected clusters to enhance resolution and improve source attribution accuracy; and (3) **Genetic validation**—recombination analysis to assess population-level genetic phenomena. Tier 1-2 analysis focuses exclusively on algorithmic resolution validation, intentionally excluding recombination considerations to evaluate the algorithm’s core clustering performance, while Tier 3 provides population genetic context to assess potential confounding factors [Pightling et al., 2018b, Taylor et al., 2015].

**Tier 1: Global Resolution Assessment via Single Linkage Clustering**

We first assessed cgDist’s impact on global clustering patterns using single linkage hierarchical clustering as a standardized analytical tool. Single linkage clustering was performed using Python 3.11 with SciPy v1.11.4 on all four datasets (Lm-1426, Lm-1874, Se-1540, Se-1434) comparing cgMLST distance against both cgDist modes (SNPs-only and SNPs+InDel-events). For clustering congruence analysis using the Adjusted Rand Index, we evaluated a range of thresholds (1-15 alleles) to assess method agreement across different stringency levels. This method provides a systematic framework to evaluate how enhanced genomic resolution affects cluster detection at the population level.

The global analysis examines three critical aspects: (1) **Cluster Resolution Enhancement**—measuring how cgDist increases the number of detected clusters and changes cluster size distributions compared to cgMLST distance; (2) **Cluster Composition Changes**—quantifying which isolates change cluster assignments and the epidemiological impact of reassignments; and (3) **Clustering Congruence**—evaluating consistency between methods using the Adjusted Rand Index to quantify the proportion of samples experiencing cluster assignment changes.

Hierarchical clustering using the single linkage method was performed on cgMLST distance and cgDist matrices (both SNPs-only and SNPs+InDel events modes) at epidemiologically critical thresholds (≤7 alleles for *L. monocytogenes* [Moura et al., 2016], ≤10 alleles for *S. enterica* [European Food Safety Authority, 2022, Radomski et al., 2019, Ruppitsch et al., 2015b, Alikhan et al., 2018]). All four datasets were filtered to 98% sample completeness threshold (*θ*_*s*_ = 0.98) [Katz et al., 2017, Carroll et al., 2021].

The analysis quantifies false association prevention—the number of isolate pairs that would be incorrectly clustered together using cgMLST distance but are appropriately separated using cgDist nucleotide-level analysis [Chen et al., 2020, Allard et al., 2016]. This metric directly measures the epidemiological value of enhanced genomic resolution by counting prevented false epidemiological associations that could mislead outbreak investigations [Gymoese et al., 2017, Ruppitsch et al., 2018]. Additionally, we analyze cluster splitting patterns to identify large cgMLST distance clusters that are subdivided into epidemiologically meaningful subclusters by cgDist, enabling more precise source attribution and outbreak scope determination [Moura et al., 2017, Kleta et al., 2017].

**Tier 2: Food Safety Source Prioritization via Star Clustering**

Following global assessment, we implemented patient-centered source prioritization analysis using star clustering topology to demonstrate cgDist’s optimal application in food safety investigations. Star clustering, the standard method employed by regulatory agencies including EFSA for outbreak investigation [European Food Safety Authority (EFSA), 2024], provides the ideal framework for evaluating cgDist’s “cluster zoom” capability within epidemiologically relevant outbreak scenarios.

Star clustering reflects the epidemiological reality of food safety investigations, where clinical isolates (patients) serve as known transmission endpoints and investigators trace backwards to identify potential sources [Brown et al., 2019, Kleta et al., 2017]. In star clustering, all isolates must be within the specified threshold distance directly from the clinical isolate, creating well-defined sets of closely related isolates around each patient case and enabling unambiguous source prioritization based solely on patient-source genetic distances.

For each clinical isolate (patient), we constructed star clusters using Python 3.11 and NetworkX v3.2 [Pightling et al., 2018b, Allard et al., 2018] where the patient serves as the cluster center and all non-patient isolates (food, environmental, processing plant samples) within established epidemiological thresholds serve as cluster members. Only patients with at least one source within the epidemiological threshold can form clusters. Within each star cluster, potential sources are ranked by their genomic distance to the patient isolate using hierarchical tiebreaking: primary sorting by cgDist distance, secondary sorting by cgMLST distance for tied cgDist values, and tertiary lexicographic sorting by sample identifier for remaining ties. This hierarchical approach ensures that cgDist rankings preserve the fundamental epidemiological relationships established by cgMLST while adding nucleotide-level granularity, enabling prioritization of investigation efforts with lower-ranked sources receiving higher investigative priority.

The analysis utilized the 1,426 *L. monocytogenes* and 1,540 *S. enterica* datasets with comprehensive source classification metadata essential for distinguishing patients from potential food sources. Critical rank shifts are defined as patient-source relationships where sources ranked within the top 3 positions by cgMLST distance are displaced outside the top 3 positions by cgDist analysis, representing the most significant changes in source prioritization that could impact investigation focus and resource allocation [Allard et al., 2018, Pightling et al., 2018b].

**Tier 3: Integrated Recombination Analysis**

Following global screening and source prioritization analysis, cgDist’s unified cache architecture enables integrated recombination analysis by leveraging alignment statistics computed during standard distance calculations. When caches are enriched with sequence length information (an optional user-configurable setting that increases cache size, with the trade-off varying by species based on genome size and allelic diversity), the system can identify potential recombination events through mutation density analysis 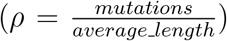 without additional computational overhead [Arnold et al., 2018, Didelot and Maiden, 2010, Feil et al., 2003, Croucher et al., 2015]. Depending on the organism’s population structure, users can optimize cache size: for strictly clonal species with negligible recombination, enrichment is less critical, whereas for panmictic species where recombination is frequent, appropriate enrichment helps capture the genetic di-versity necessary for accurate distance calculations [Smith et al., 1993, Didelot et al., 2010].

The algorithm applies an epidemiological filtering threshold (Hamming distance ≤15 allelic differences) to restrict recombination analysis to sample pairs within plausible epidemiological relationships. This threshold extends the canonical ≤10-allele outbreak detection threshold [Radomski et al., 2019, European Food Safety Authority, 2022] to include moderately divergent samples that may still harbor epidemiologically relevant recombination signals, while excluding highly divergent pairs unlikely to represent recent transmission events. A 3.0% mutation density threshold identifies sequences exceeding recombination thresholds. The Se-1540 *S. enterica* dataset was selected for calibration due to its substantial recombination activity, in contrast to *Listeria monocytogenes*, which exhibits significantly lower recombination rates, as supported by studies reporting high recombination in *Salmonella* [Diez-Villasenor and Rodriguez-Valera, 2021, *Gonzalez-Escalona et al*., *2021] and low recombination in Listeria* [Orsi et al., 2008, Cai et al., 2008]. The algorithm supports optional loci filtering, and all parameters are configurable to accommodate different epidemiological characteristics and recombination patterns of various pathogens.

**Performance Analysis and Incremental Processing** We conducted comprehensive scalability analysis using high-performance computing infrastructure and developed a Nextflow pipeline for incremental processing validation. The pipeline simulates temporal arrival of isolates in surveillance scenarios, measuring computational performance, cache efficiency, and alignment reuse benefits as surveillance programs continuously integrate new genomic data. Detailed methodology for performance benchmarking, threading analysis, and the complete Nextflow pipeline architecture are described in Supplementary Methods Sections 7 and 7.

## 3 Results

### 3.1 Mathematical Validity and Consistency

#### 3.1.1 Mathematical Constraint Validation

Validation confirmed that all distance calculations satisfied the fundamental design constraint established by the cgMLST-cgDist Ordering Theorem that `cgDist` values are greater than or equal to corresponding cgMLST distances: (*d*_cgMLST_ ≤ *d*_cgDist_). Comprehensive testing across all four datasets totaling 4,983,517 pairwise comparisons confirmed zero violations of the expected ordering relationships: 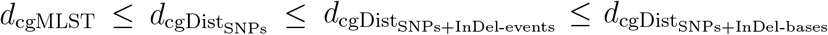. This comprehensive testing validates both the theoretical algorithm design and practical implementation accuracy across diverse bacte-rial species, dataset sizes, and genetic diversity levels (Table 1).

**Table 1:** Correlation Analysis Between Distance Methods. All correlations are significant at p *<* 0.001. r = Pearson correlation coefficient, *ρ* = Spearman rank correlation coefficient.

| Comparison | Lm-1426 | Lm-1874 | Se-1540 | Se-1434 |
| --- | --- | --- | --- | --- |
| | (1,067,991 pairs)<br>$r / \rho$ | (1,755,001 pairs)<br>$r / \rho$ | (1,185,030 pairs)<br>$r / \rho$ | (1,027,461 pairs)<br>$r / \rho$ |
| SNPs vs cgMLST | 0.771 / 0.933 | 0.788 / 0.876 | 0.716 / 0.931 | 0.901 / 0.905 |
| SNPs+InDel-events vs cgMLST | 0.771 / 0.933 | 0.789 / 0.880 | 0.716 / 0.931 | 0.901 / 0.905 |
| SNPs+InDel-events vs SNPs | 1.000 / 1.000 | 1.000 / 1.000 | 1.000 / 1.000 | 1.000 / 1.000 |
| SNPs+InDel-bases vs cgMLST | 0.771 / 0.921 | 0.789 / 0.872 | 0.699 / 0.890 | 0.772 / 0.784 |
| SNPs+InDel-bases vs SNPs | 1.000 / 0.994 | 1.000 / 0.996 | 0.966 / 0.947 | 0.891 / 0.911 |
| SNPs+InDel-bases vs SNPs+InDel-events | 1.000 / 0.994 | 1.000 / 0.996 | 0.966 / 0.949 | 0.893 / 0.914 |

**Note:** All datasets: 100% completeness threshold. Lm-1426, Lm-1874: *L. monocytogenes*. Se-1540, Se-1434: *S. enterica*.

Strong correlations between distance methods confirmed algorithmic consistency (Pearson 0.699-1.000, Spearman 0.784-1.000, all p *<* 0.001). Perfect correlations between SNPs+InDel-events and SNPs modes reflect the additive nature of InDel events, while high correlations with cgMLST distances demonstrate that cgDist preserves fundamental epidemiological relationships while adding nucleotide-level granularity.

### 3.2 Multi-scale Clustering Analysis: From Global Screening to Cluster Expansion

#### 3.2.1 Tier 1: Global Resolution Assessment via Single Linkage Clustering

Single linkage clustering analysis on complete 98% filtered datasets reveals substantial variation in maximum cluster sizes and demonstrates how cgDist systematically subdivides large clusters to prevent false outbreak associations (Table 2). Maximum cluster sizes vary significantly across datasets: Lm-1426 exhibits the largest clusters (up to 201 samples with Hamming distance, 199 with cgDist methods), while Se-1434 shows more conservative clustering (maximum 31 samples with Hamming, 26-28 with cgDist). Lm-1874 and Se-1540 demonstrate intermediate patterns with maximum cluster sizes of 43 and 124-125 samples respectively.

**Table 2:** Largest cluster subdivisions by cgDist methods across all datasets using single linkage clustering. Single linkage clusters identified by cgMLST distance at epidemiological thresholds (≤7 for *Listeria*, ≤ 10 for *Salmonella*) are subdivided by cgDist into distinct genetic groups. Rankings show the 5 largest original cgMLST clusters (by size) that were subdivided by cgDist analysis for each dataset and cgDist mode. cgDist Subdivision uses bracket notation for multi-sample clusters with inter-cluster distances followed by singleton count (e.g., “[5,2](d*≥*12)+4*×*1” indicates two clusters of 5 and 2 samples separated by *≥*12 distance, plus 4 singletons).

| Dataset | cgDist Mode | cgMLST Rank | cgMLST Size | cgDist Groups | cgDist Subdivision |
| --- | --- | --- | --- | --- | --- |
| Lm-1426 | SNPs-only | 1 | 201 | 3 | 199+2×1 |
| | | 2 | 169 | 3 | 167+2×1 |
| | | 3 | 34 | 2 | 33+1×1 |
| | SNPs+InDel-events | 1 | 201 | 3 | 199+2×1 |
| | | 2 | 169 | 4 | 166+3×1 |
| | | 3 | 44 | 2 | 43+1×1 |
| | | 4 | 34 | 2 | 33+1×1 |
| | | 5 | 17 | 3 | 15+2×1 |
| Lm-1874 | SNPs-only | 1 | 39 | 2 | 38+1×1 |
| | SNPs+InDel-events | 1 | 39 | 2 | 38+1×1 |
| | | 2 | 11 | 2 | [6,5]($d \geq 8$) |
| Se-1540 | SNPs-only | 1 | 125 | 2 | 124+1×1 |
| | | 2 | 122 | 6 | 117+5×1 |
| | | 3 | 75 | 6 | 70+5×1 |
| | | 4 | 72 | 3 | 70+2×1 |
| | | 5 | 44 | 3 | 42+2×1 |
| | SNPs+InDel-events | 1 | 125 | 2 | 124+1×1 |
| | | 2 | 122 | 7 | 116+6×1 |
| | | 3 | 75 | 11 | 65+10×1 |
| | | 4 | 72 | 6 | 67+5×1 |
| | | 5 | 44 | 4 | 41+3×1 |
| Se-1434 | SNPs-only | 1 | 31 | 4 | 28+3×1 |
| | | 2 | 30 | 4 | 27+3×1 |
| | | 3 | 18 | 2 | 17+1×1 |
| | | 4 | 15 | 2 | 14+1×1 |
| | | 5 | 12 | 6 | [6,2]($d \geq 11$)+4×1 |
| | SNPs+InDel-events | 1 | 31 | 7 | 25+6×1 |
| | | 2 | 30 | 5 | 26+4×1 |
| | | 3 | 18 | 2 | 17+1×1 |
| | | 4 | 15 | 2 | 14+1×1 |
| | | 5 | 12 | 6 | [6,2]($d \geq 11$)+4×1 |

**Note:** 98% completeness datasets. Lm-1426 (n=1,381), Lm-1874 (n=1,824): *L. monocytogenes*. Se-1540 (n=1,446), Se-1434 (n=1,192): *S. enterica*. Shows clusters $\geq 10$ samples subdividing into multiple groups.

**Species-Specific Subdivision Patterns** Analysis reveals distinct patterns between bacterial species. *Listeria monocytogenes* demonstrates conservative subdivision: the largest cluster in Lm-1426 (201 samples, all within ≤7 alleles) subdivides into 199 closely related samples plus 2 genetic outliers exceeding *≥*8 SNPs from the main group, potentially representing contamination, mislabeling, or convergent evolution. The 169-sample cluster similarly shows minimal fragmentation. Lm-1874 provides evidence of differential resolution between cgDist modes, with an 11-sample cluster remaining intact under SNPs-only analysis but subdividing into two distinct groups of 6 and 5 samples (separated by ≥ 8 SNPs) under SNPs+InDel-events analysis. *Salmonella enterica* exhibits more extensive fragmentation reflecting higher genetic diversity: a 75-sample cluster in Se-1540 subdivides into 11 distinct genetic groups (65 main + 10 singletons), suggesting multiple introduction events incorrectly merged by allelic similarity. Se-1434 demonstrates true multi-cluster subdivision where clusters split into multiple subclusters (e.g., one 8-sample cluster splitting into subclusters of 6 and 2 samples separated by *≥*11 SNPs, plus 4 singleton outliers), indicating genetic subpopulations separated by epidemiologically significant distances.

**Clustering Congruence:** Despite resolution enhancement, clustering methods showed high congruence as measured by Adjusted Rand Index (ARI) values at epidemiological thresholds (Table 3, Figure 3). Lm-1426 and Lm-1874 datasets achieved ARI values of 0.842-1.000, indicating that cgDist enhances resolution while preserving most existing cluster relationships. Se-1540 and Se-1434 datasets showed lower but still substantial congruence (ARI 0.553-1.000), reflecting the greater restructuring potential in these more genetically diverse datasets. These ARI values demonstrate that cgDist provides enhanced discrimination without fundamentally disrupting epidemiologically valid associations established by cgMLST distance clustering.

**Table 3:**
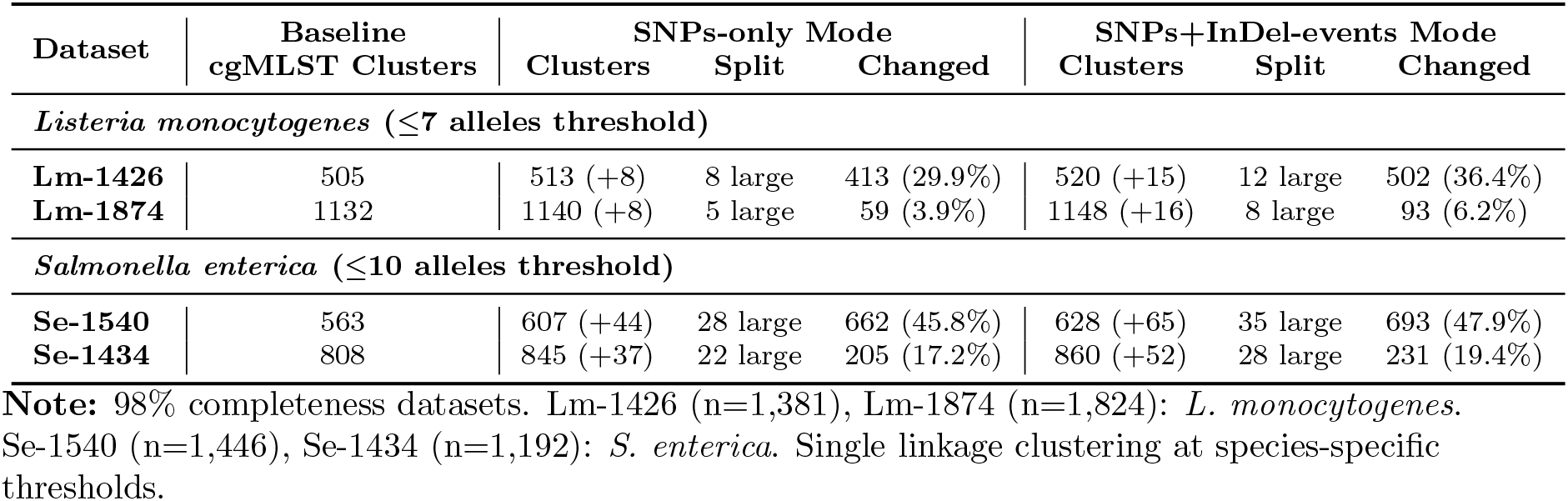
Concrete Clustering Changes: How cgDist Affects Outbreak Cluster Detection using single linkage clustering. Numbers in parentheses show additional clusters compared to cgMLST distance baseline. Split values indicate the number of large cgMLST distance clusters (*≥* 10 samples) identified by single linkage clustering that were subdivided by cgDist analysis. Changed values show the number and percentage of samples that experienced cluster reassignment when using cgDist instead of cgMLST distance.

**Figure 3.**
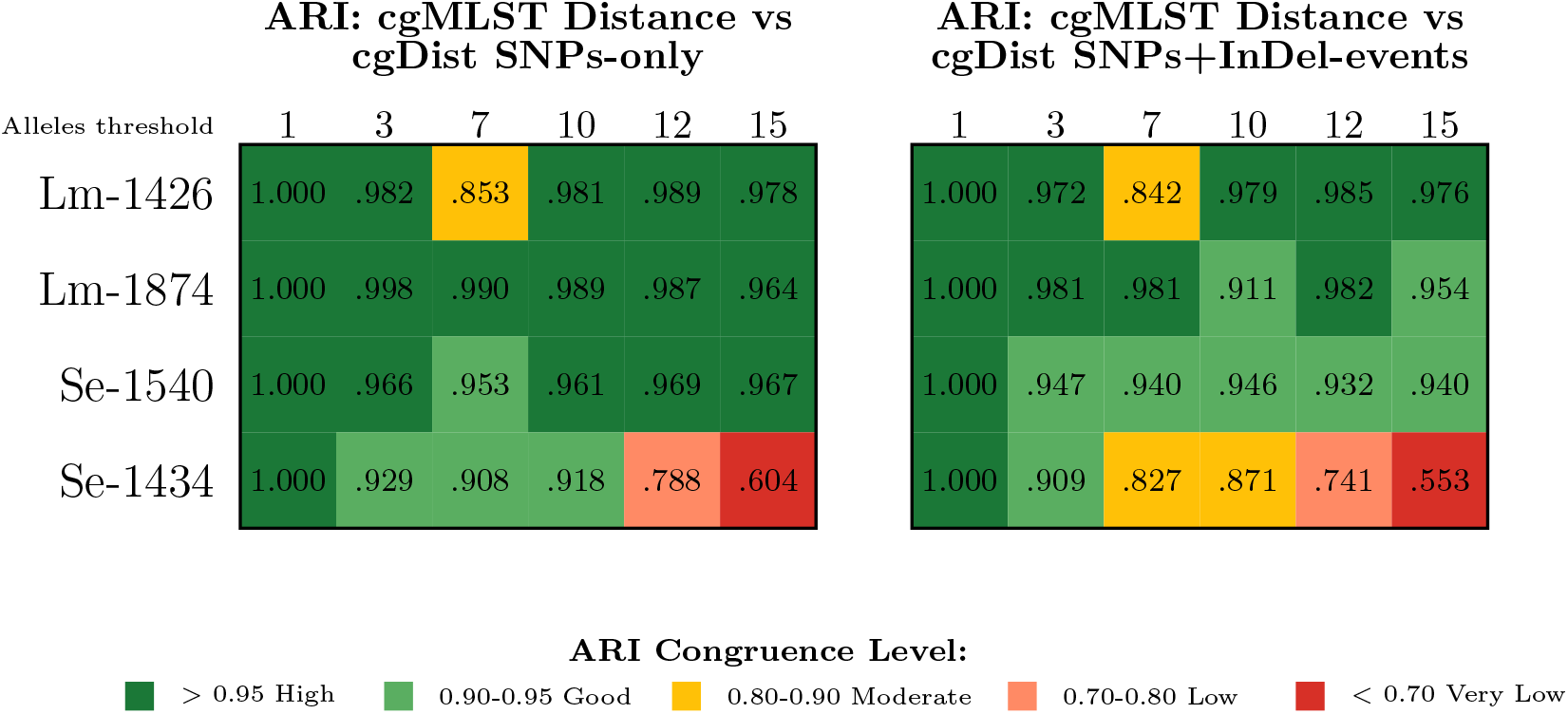
Threshold-Dependent ARI Congruence Analysis Between cgMLST Distance and cgDist Modes using single linkage clustering. Heatmaps display Adjusted Rand Index values measuring congruence between single linkage clustering results at selected critical thresholds (1, 3, 7, 10, 12, 15 alleles). Each cell represents ARI between traditional cgMLST distance single linkage clustering and enhanced cgDist single linkage clustering at the specified threshold. Color intensity indicates congruence level: dark green (high congruence, ARI *>* 0.95), medium green (good congruence, 0.90-0.95), yellow (moderate changes, 0.80-0.90), orange (low congruence, 0.70-0.80), red (substantial restructuring, *<* 0.70). Left panel compares cgMLST distance vs SNPs-only mode; right panel compares cgMLST distance vs SNPs+InDel-events mode. *Listeria monocytogenes* demonstrates consistently high congruence across thresholds, while *Salmonella enterica* Se-1434 exhibits progressive degradation culminating in substantial restructuring at higher thresholds (ARI 0.553-0.604).

**Clustering Composition Changes:** Clustering composition analysis reveals the concrete impact of enhanced genomic resolution on epidemiological cluster detection across all validation datasets (Table 3). cgDist changes how outbreak clusters are detected and defined, with dataset-specific patterns reflecting the genetic diversity present in each sample collection. For *L. monocytogenes*, cgDist provides enhancement that preserves most existing epidemiological associations while refining ambiguous cases. The Lm-1426 dataset shows substantial clustering changes (29.9% samples in SNPs-only, 36.4% in SNPs+InDel-events mode), while Lm-1874 demonstrates more conservative changes (3.9% and 6.2% respectively), indicating dataset-specific genetic diversity impacts. For *S. enterica*, cgDist reveals hidden diversity within cgMLST distance clusters, reflecting the species’ higher genetic diversity. The Se-1540 dataset demonstrates significant subdivision potential with 45.8% samples changing assignments in SNPs-only mode and 47.9% in SNPs+InDel-events mode. The Se-1434 dataset shows more moderate changes (17.2% and 19.4% respectively), demonstrating variable genetic diversity across datasets. The systematic increase in cluster numbers, combined with dataset-specific change patterns, indicates that cgDist primarily splits existing clusters rather than randomly reassigning isolates, targeting epidemiologically ambiguous relationships where nucleotide-level analysis provides clearer discrimination.

#### 3.2.2 Tier 2: Food Safety Source Prioritization via Star Clustering

Our multi-scale analysis framework demonstrates cgDist’s application: initial cluster identification using single linkage clustering followed by targeted source prioritization using star clustering methodology. Star clustering analysis, the regulatory standard for outbreak investigation employed by agencies such as EFSA [European Food Safety Authority (EFSA), 2024], provides a framework for evaluating cgDist’s enhanced resolution within epidemiologically important outbreak scenarios.

**Star Clustering Implementation:** Initial screening using cgMLST distance on quality-filtered datasets (98% completeness threshold) identified 264 *Listeria* clusters from 1,070 total clinical samples (Lm-1426 dataset) and 136 *Salmonella* clusters from 847 total clinical samples (Se-1540 dataset). These clusters represent the starting point for multi-scale analysis—each cluster becomes a focused unit where cgDist expands internal resolution to reveal hidden genetic relationships (Table 4).

**Table 4:** Median cluster sizes across distance calculation methods using star clustering methodology on 98% completeness-filtered datasets. Star clustering performed using patient-centered methodology with epidemiological thresholds of ≤7 alleles for *Listeria* and ≤10 alleles for *Salmonella*. cgMLST distance clustering establishes the foundational topology, while cgDist methods (SNPs-only and SNPs+InDel-events) are used for source prioritization analysis within established clusters.

| Dataset | Clinical<br>Samples | Median Cluster Size (Range) |  |  |
| --- | --- | --- | --- | --- |
|  |  | cgMLST Distance | cgDist SNPs-only | cgDist SNPs+InDel-events |
| Lm-1426 | 1,070 | 2 (2–91) | 2 (2–89) | 2 (2–87) |
| Se-1540 | 847 | 34 (2–100) | 54 (2–89) | 68 (2–87) |

**Note:** 98% completeness datasets. Star clustering analysis limited to datasets with comprehensive source attribution metadata (Lm-1426 and Se-1540). Other datasets (Lm-1874, Se-1434) lack detailed source classification required for patient-centered clustering methodology.

Within each identified cluster, cgDist’s enhanced resolution expands our understand-ing of source-patient relationships, revealing priority reorderings invisible at the allelic level (Table 5). *L. monocytogenes* clusters show moderate internal reorganization from SNPs-only (25.8% clusters affected) to SNPs+InDel-events mode (31.1% clusters affected), demonstrating that even within small, tight clusters, cgDist expands resolution to differentiate closely related sources. Remarkably, despite this internal expansion, top source stability is maintained with no rank 1 changes, indicating that cgDist refines rather than disrupts primary epidemiological associations.

**Table 5:** Priority reordering impact across cgDist modes for datasets with source attribution metadata. Priority changes calculated between cgMLST distance and cgDist rankings using hierarchical tiebreaking (SNPs → cgMLST → Sample ID). Critical rank shifts: sources ranked 1-3 by cgMLST distance displaced to *>* 3 by cgDist (loss of top priority). Top 3 changes: sources remaining in top 3 but changing position.

| Species | Metric | cgDist Mode |  |
| --- | --- | --- | --- |
|  |  | SNPs-only | SNPs+InDels |
| *L. monocytogenes* | Patients with sources | 264 | 264 |
|  | Clusters with reordering | 68 (25.8%) | 82 (31.1%) |
|  | Rank 1 changes | 0 (0.0%) | 0 (0.0%) |
|  | Top 3 changes | 1 (0.4%) | 2 (0.8%) |
|  | Top 10 changes | 17 (6.4%) | 26 (9.8%) |
|  | Average rank shift | 3.39 | 3.60 |
|  | Maximum rank shift | 74 | 75 |
| | Most extreme shift | Lm_0114: 15→89 | Lm_0114: 15→90 |
|  | Critical rank shifts | 1 | 2 |
| *S. enterica* | Patients with sources | 136 | 136 |
|  | Clusters with reordering | 76 (55.9%) | 82 (60.3%) |
|  | Rank 1 changes | 4 (2.9%) | 4 (2.9%) |
|  | Top 3 changes | 16 (11.8%) | 16 (11.8%) |
|  | Top 10 changes | 41 (30.1%) | 43 (31.6%) |
|  | Average rank shift | 6.66 | 6.99 |
|  | Maximum rank shift | 81 | 81 |
| | Most extreme shift | Se_0439: 87→6 | Se_0439: 87→6 |
|  | Critical rank shifts | 16 | 16 |

**Note:** 98% completeness datasets. Analysis limited to datasets with comprehensive source attribution metadata: Lm-1426 (*L. monocytogenes*) and Se-1540 (*S. enterica*). Thresholds: $\leq 7$ alleles (*Listeria*), $\leq 10$ (*Salmonella*).

*S. enterica* clusters exhibit more extensive internal restructuring from SNPs-only (55.9% clusters affected) to SNPs+InDel-events mode (60.3% clusters affected). This greater sensitivity to cluster expansion reflects the species’ higher genetic diversity, where cgDist reveals substantial hidden variation within cgMLST-defined clusters. The consistent moderate top source changes (2.9% rank 1 changes in both modes) demonstrate that cluster expansion can fundamentally alter outbreak investigation priorities.

The magnitude of priority changes revealed consistent patterns across cgDist modes with species-specific amplification effects. *Listeria* demonstrated relatively stable rank shift magnitudes between SNPs-only (average 3.39 positions, maximum 74) and SNPs+InDel-events modes (average 3.60 positions, maximum 75), indicating minimal but measurable additive impact from InDel events on source prioritization. *Salmonella* showed comparable stability between modes, with SNPs-only (average 6.66 positions, maximum 81) and SNPs+InDel-events (average 6.99 positions, maximum 81) exhibiting similar reordering magnitudes, suggesting that species-specific genetic diversity patterns, rather than InDel inclusion, primarily drive prioritization differences. The most substantial reordering involved patient Se 0003 (Figure 6), where 11 sources were completely tied at 1 allele difference by cgMLST. cgDist revealed that source Se 0695 had only 1 SNP while other tied sources had 2-7 SNPs, correctly identifying it as the biologically closest source (rank 1). Simultaneously, sources initially ranked 1-2 by cgMLST were demoted to ranks 9-11, having 6-7 SNPs despite identical allelic distance, demonstrating how cgDist reveals hidden nucleotide-level variation that fundamentally alters source prioritization even among apparently equivalent candidates.

Critical rank shifts analysis revealed consistent patterns across cgDist modes, with species-specific thresholds for regulatory significance (Figure 4). *Listeria* showed minimal critical displacement increases from SNPs-only (1 critical shift) to SNPs+InDel-events mode (2 critical shifts), maintaining relatively stable top-priority source identification. *Salmonella* demonstrated identical critical shift counts across both modes (16 critical shifts each), indicating that primary priority reordering occurs at the SNPs level rather than through InDel event inclusion. These findings suggest that while InDel events provide additional granularity for distance calculations, the fundamental regulatory priorities for food safety investigations are established primarily through SNPs-level differences, with species-specific sensitivity patterns determining the magnitude of investigative impact.

**Figure 4.**
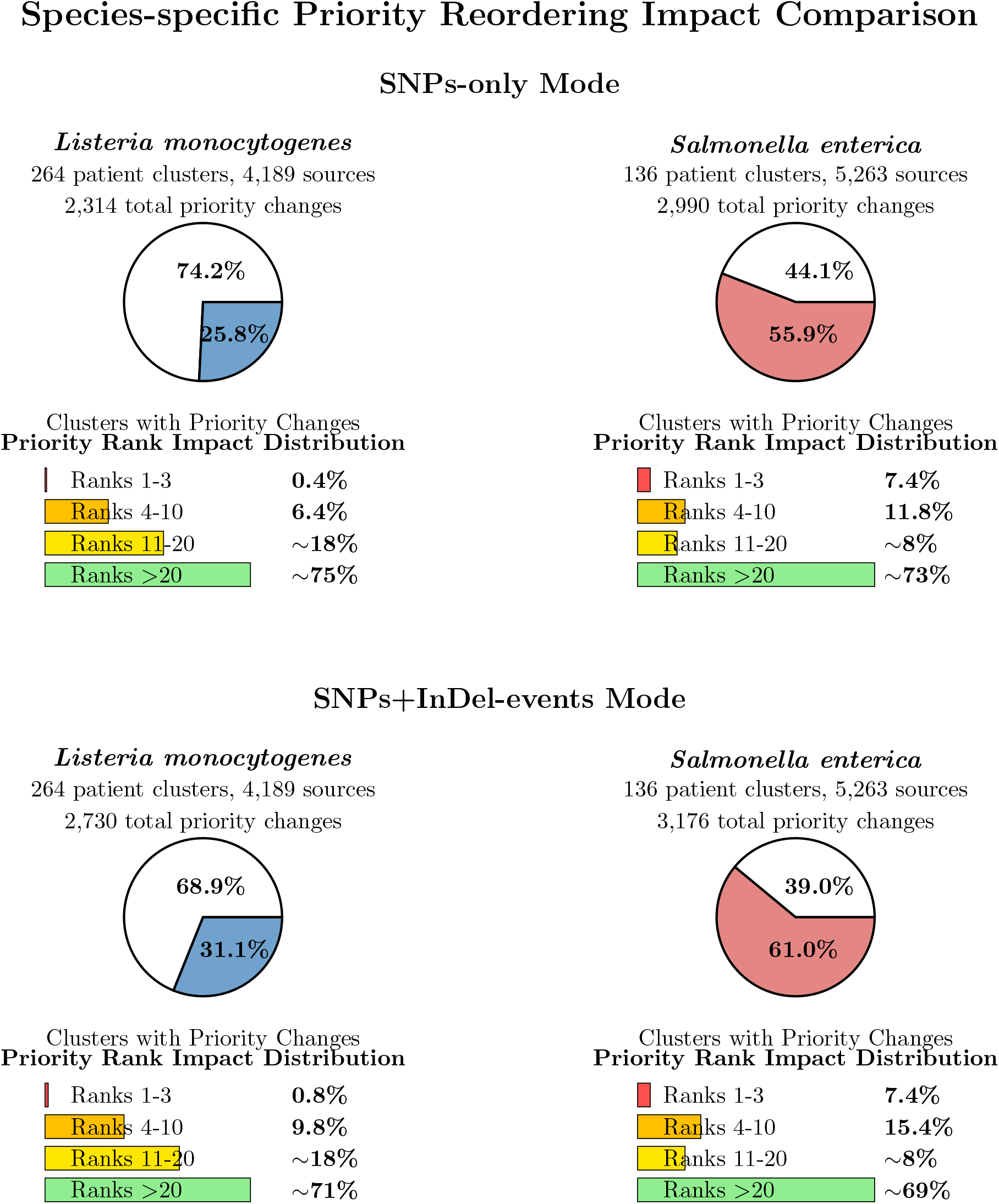
Direct comparison of species-specific source priority reordering impact between cgDist modes. Top panel shows SNPs-only mode analysis, bottom panel shows SNPs+InDel-events mode analysis. *Listeria monocytogenes* demonstrates moderate reordering increases from 25.8% to 31.1% of clusters affected, with minimal top-3 priority impact (0.4% to 0.8%). *Salmonella enterica* exhibits substantial reordering in both modes (55.9% to 61.0%) with consistent high top-3 priority impact (7.4% both modes). Species-specific sensitivity patterns to InDel event inclusion are clearly visible through vertical comparison.

**Prevention of False Source Attribution:** Comparative analysis of cgDist modes revealed systematic patterns in preventing false source attribution across both SNPs-only and SNPs+InDel-events calculation methods when compared to cgMLST distance methods (Figures 5 and 6).

**Figure 5.**
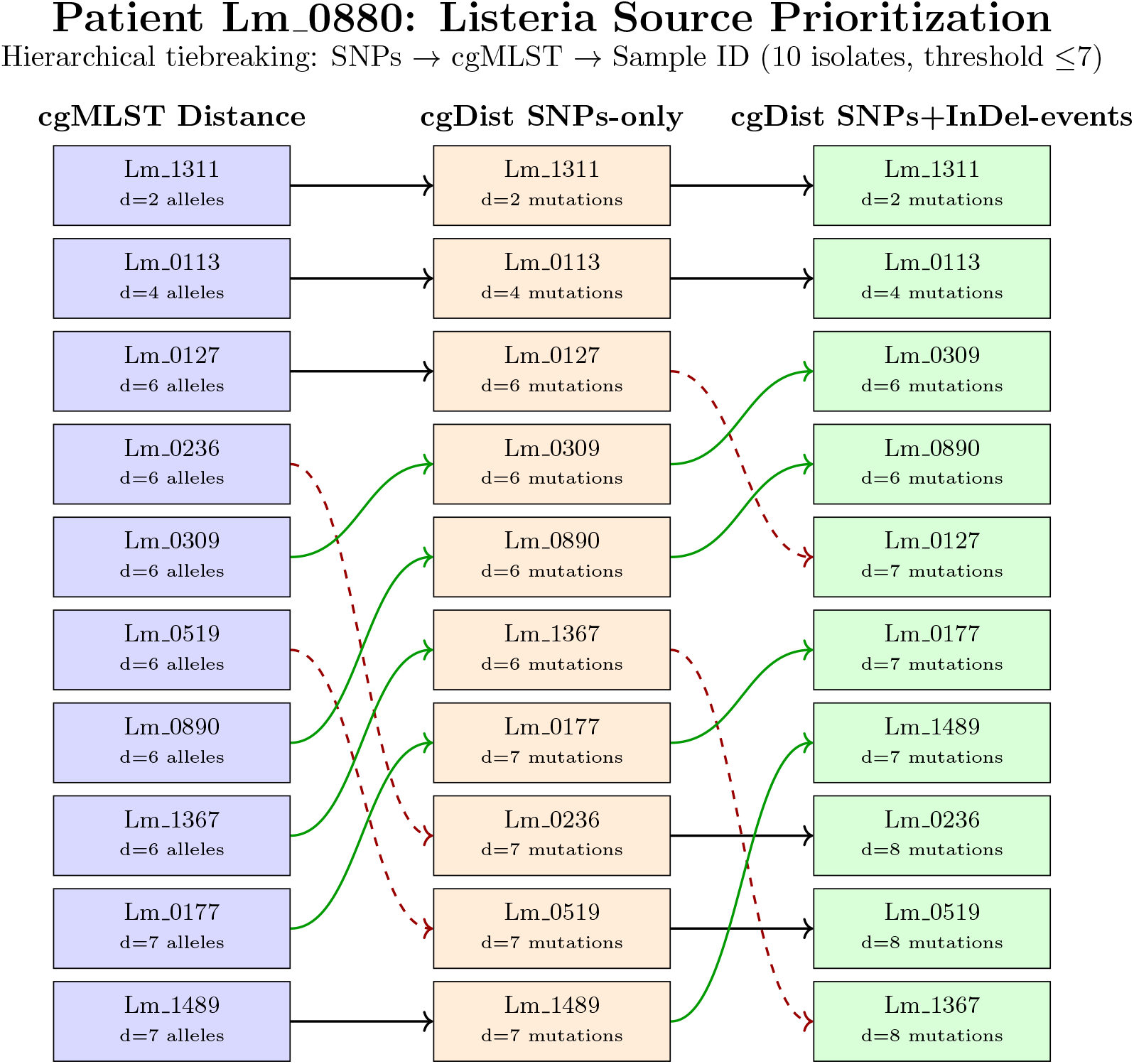
*Listeria monocytogenes* source prioritization comparing cgMLST distance versus both cgDist modes using hierarchical tiebreaking (SNPs → cgMLST → Sample ID). Patient Lm 0880 demonstrates substantial reordering between methods, with notable rank changes from cgMLST distance to SNPs-only and further movements to SNPs+InDel-events, illustrating how different mutation types contribute distinct information for *Listeria* source attribution.

**Figure 6.**
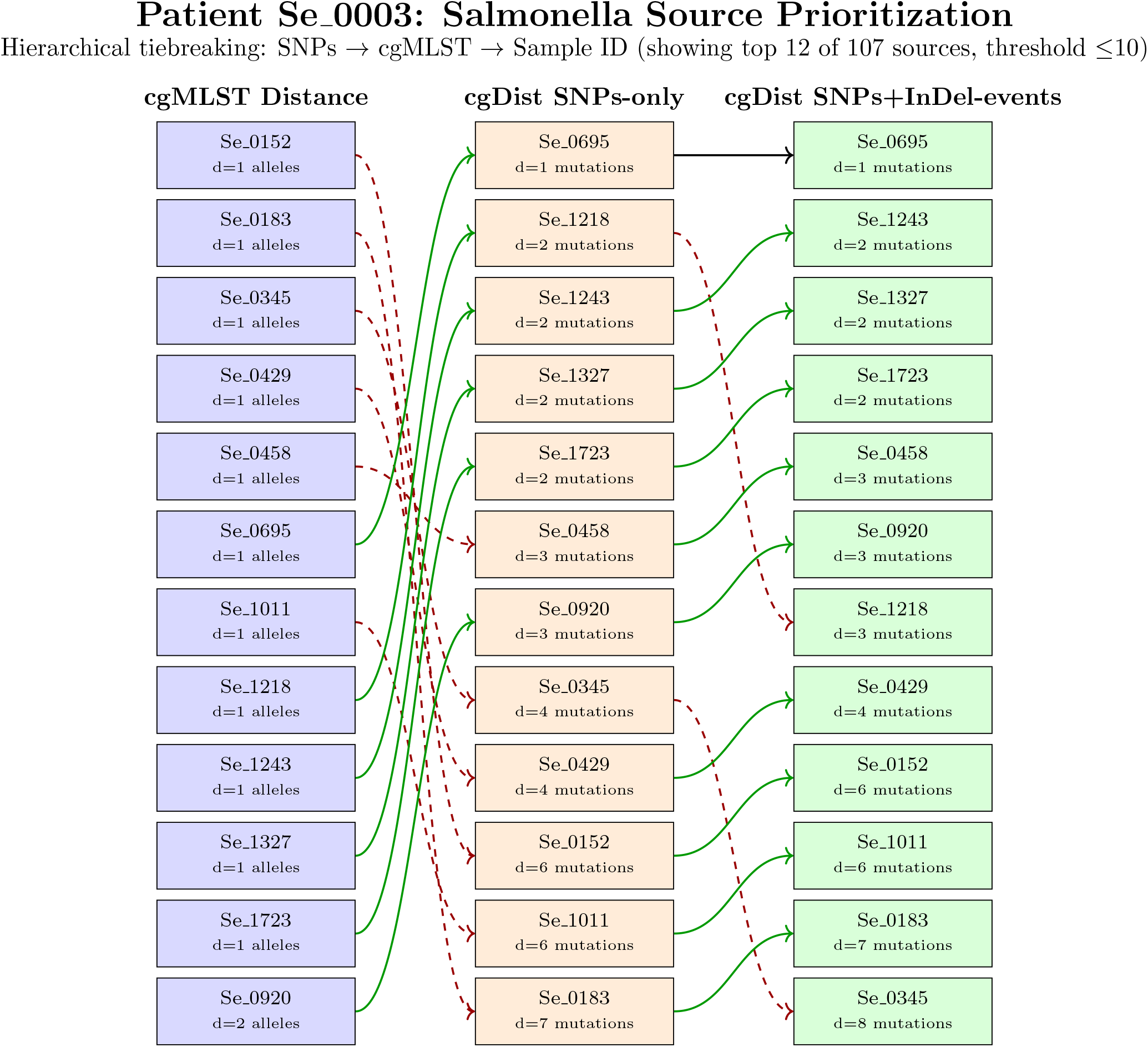
*Salmonella enterica* source prioritization comparing cgMLST distance versus both cgDist modes using hierarchical tiebreaking (SNPs → cgMLST → Sample ID). Patient Se 0003 (107 total sources) demonstrates extensive reordering between methods, with source Se 0695 advancing from rank 6 to rank 1 in both cgDist modes, while multiple highly-ranked cgMLST distance sources are substantially demoted, illustrating how different mutation types provide distinct discriminatory information for *Salmonella* source attribution.

Species-specific false attribution prevention patterns emerged from the comparative analysis. *L. monocytogenes* demonstrated conservative but consistent improvements across both cgDist modes, with minimal critical rank shifts in SNPs-only mode (1 patient-source relationship, 0.4% of clusters) increasing to modest levels in SNPs+InDel-events mode (2 patient-source relationships, 0.8% of clusters). The relatively homogeneous genetic structure of *Listeria* populations results in subtle but meaningful priority adjustments rather than extensive rank changes, indicating that both cgDist modes provide enhanced discrimination while maintaining epidemiological coherence. The incremental improvement from SNPs-only to SNPs+InDel-events mode suggests that InDel events contribute additional discriminatory power for source attribution in *Listeria* investigations.

*S. enterica* revealed more substantial false attribution prevention across both cgDist modes, reflecting the species’ higher genetic diversity. Both SNPs-only and SNPs+InDel-events modes identified identical numbers of critical rank shifts (16 patient-source relationships each, 11.8% of clusters), indicating that SNPs-level analysis provides the primary discriminatory improvement for *Salmonella* source attribution. The consistency between modes demonstrates that nucleotide-level analysis fundamentally enhances source attribution accuracy in high-diversity bacterial populations, with SNPs contributing the majority of discriminatory power and InDel events providing supplementary refinement.

Cross-mode analysis revealed that cgDist’s false attribution prevention operates through systematic enhancement of genetic relationship resolution rather than isolated correc-tions. Both cgDist modes consistently identify sources that appear epidemiologically relevant by cgMLST distance but represent distant genetic relationships upon nucleotide-level analysis. The most critical prevention scenarios involve sources initially ranked within top 3 positions by cgMLST distance but displaced outside epidemiological relevance by cgDist analysis, enabling more accurate resource allocation during food safety investigations and reducing the risk of misdirected regulatory actions.

#### 3.2.3 Tier 3: Recombination Analysis

cgDist’s integrated recombination analysis demonstrates the capability to support multiscale surveillance workflows through zero-overhead cache reuse, completing the three-tier analytical framework. By leveraging enriched cache data containing sequence length information, the system enables mutation density analysis 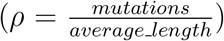 to identify potential horizontal gene transfer events without additional computational overhead.

**Empirical Threshold Determination:** Analysis of mutation density distribution in the Se-1540 *S. enterica* dataset revealed a bimodal distribution with modes at 4.9% and 16.5%, median density 3.50%, and first quartile 2.90%. The empirically-determined 3.0% mutation density threshold was positioned near the median and closely aligned with natural population structure, capturing 72.6% of recombination events while maintaining conservative specificity and effectively discriminating between typical allelic variation and genuine horizontal gene transfer.

**Recombination Detection Results:** Using this 3.0% threshold, we identified recombination events in 20 sample pairs in the Se-1540 dataset affecting 0.03% of EFSA loci. Notably, all samples involved in recombination events with available metadata were patient samples, while most lacked temporal information, contrasting with 50% of the dataset that includes collection years spanning 1987-2021. Analysis across all 98% filtered datasets revealed striking species-specific differences in recombination activity. Only the *S. enterica* Se-1540 dataset exhibited detectable horizontal gene transfer events exceeding the 3.0% mutation density threshold. In contrast, the *L. monocytogenes* datasets (Lm-1426, Lm-1874) and the second *S. enterica* dataset (Se-1434) showed minimal recombination activity (1 sample pair), indicating predominantly clonal inheritance patterns. This species- and population-specific variation has important implications for genomic surveillance strategies and outbreak investigation protocols.

**Clustering Robustness Validation:** Recombination analysis revealed differential patterns across datasets with important implications for our Tier 1-2 analytical framework. Se-1540 showed 20 sample pairs exhibiting recombination events affecting 0.03% of EFSA loci, while Se-1434 demonstrated minimal recombination (1 pair). Metadata analysis revealed that among samples involved in Se-1540 recombination events, all those with available temporal information lacked collection years, contrasting with 50% of the dataset that includes years spanning 1987-2021. Additionally, all samples involved in recombination events with known metadata were patient samples, representing potential clinical infections across Netherlands (2012) and Portugal (2017). Critically, these recombination events occurred primarily between samples that were already separated in our clustering analysis, indicating that distance-based clustering naturally segregates samples with extensive horizontal gene transfer from epidemiologically relevant transmission clusters. Since patient samples serve as cluster centers in star clustering analysis rather than potential sources for ranking, these recombination events do not affect source prioritization rankings. This demonstrates that our Tier 1-2 analytical framework successfully focuses on epidemiologically relevant genetic variation while maintaining robustness against population-level genetic phenomena.

Detailed recombination detection results and clustering robustness analysis are provided in Supplementary Results Section 7.

### 3.3 Performance Characteristics and Deployment Validation

cgDist demonstrates dataset-dependent scalability patterns that correlate with genetic diversity rather than dataset size, achieving optimal parallel performance at 16-32 threads for high-diversity populations and near-linear scaling for low-diversity datasets. The unified cache architecture provides substantial performance improvements for multi-mode workflows, with cache reuse delivering up to 14*×* throughput improvements and 94% time reduction across analysis modes. Comprehensive performance analysis, scalability benchmarks, and memory efficiency evaluation are detailed in Supplementary Results Section 7.

### 3.4 Surveillance Pipeline Applications

cgDist has been successfully integrated into the GENPAT bioinformatics platform [de Ruvo et al., 2024] where it serves as the core distance calculation engine for real-time epidemiological analysis. Incremental processing validation confirms that cgDist’s cache-aware design enables practical deployment of nucleotide-level genomic surveillance at the scale and speed required for modern public health practice. Detailed performance results from production surveillance integration and incremental processing validation are provided in Supplementary Results Section 7.

## 4 Discussion

### 4.1 Methodological Innovation and Validation

cgDist successfully bridges the resolution gap between traditional cgMLST and computationally intensive SNP-calling approaches while maintaining the standardization and efficiency required for routine surveillance. The comprehensive validation across 4,983,517 pairwise comparisons confirms the algorithm’s mathematical soundness, with perfect preservation of the ordering constraint *d*_cgMLST_ ≤ *d*_cgDist_ and strong correlations between distance methods (Pearson *r* 0.699-1.000, Spearman *ρ* 0.784-1.000). This consistency demonstrates that enhanced nucleotide-level resolution maintains epidemiological relevance while providing additional discriminatory power [Schürch et al., 2018, Didelot et al., 2017].

The unified cache architecture represents a significant departure from traditional approaches that precompute entire schema combinations. By processing only allele pairs present in the current dataset and storing comprehensive alignment statistics in reusable cache entries, cgDist achieves computational efficiency comparable to cgMLST while providing nucleotide-level granularity.

### 4.2 Tier 1-2 Validation: The Cluster Zoom Paradigm

Our multi-tier validation framework demonstrates cgDist’s effectiveness across different surveillance contexts, with the most significant practical contribution emerging from its ability to “zoom into” specific outbreak clusters for detailed investigation. The Tier 1-2 analysis intentionally focuses on distance-based clustering performance for outbreak investigation workflows, where enhanced SNP-level resolution provides critical epidemiological value without the complexity of population-level genetic phenomena. While global ARI values (0.553-1.000) suggest modest overall restructuring, targeted analysis within epidemiologically relevant clusters reveals substantial internal reorganization invisible to traditional methods. In star clustering analysis, 25.8-31.1% of *Listeria monocytogenes* clusters and 55.9-61.0% of *Salmonella enterica* clusters showed source priority reordering, directly impacting outbreak investigation decisions.

While cluster-focused analysis approaches exist in tools like PHYLOViZ and ReporTree that zoom from many samples to few samples [Ribeiro-Gonçalves et al., 2023, Nascimento et al., 2017], cgDist provides a complementary capability: zooming from allelic distances to nucleotide-level variants within identified clusters. This deployment strategy leverages initial cgMLST screening to identify potential outbreak clusters, followed by selective application of cgDist to expand resolution from allele counts to SNP and InDel quantification within clusters requiring detailed investigation. Representative examples demonstrate systematic rather than random improvements—*Listeria* patient Lm 0880 experienced source rank changes from 4 → 8 and 7 → 4, while *Salmonella* patient Se 0003 showed a source advancing from rank 6 → 1. Such priority reorderings represent scenarios where cgMLST analysis would misdirect investigation resources toward suboptimal targets, potentially compromising outbreak response efficiency and regulatory decision-making [Pightling et al., 2018a, European Food Safety Authority, 2022].

The species-specific patterns observed in our datasets reflect the characteristic genetic features of each bacterial species: *L. monocytogenes*’ generally clonal structure yields conservative refinements within tight clusters, while *S. enterica*’ higher genetic diversity enables detection of substantial hidden variation, with some clusters fragmenting into 11 distinct genetic groups. This differential response confirms that cgDist’s value lies in targeted resolution enhancement rather than wholesale clustering restructuring.

### 4.3 Tier 3 Validation: Population Genetic Context and Clustering Robustness

The integrated recombination analysis provides essential validation of our Tier 1-2 clustering approach by demonstrating its robustness against population-level genetic phenomena. Detection of recombination events in 20 sample pairs in Se-1540 (affecting 0.03% of EFSA loci) while maintaining clustering integrity validates that distance-based methods naturally segregate samples with extensive horizontal gene transfer from those showing conventional genetic relationships.

The analysis revealed that recombination events occurred primarily between samples that were already separated in our clustering analysis, indicating minimal overlap between recombination-affected samples and established cluster boundaries. All samples involved in recombination events with available metadata were patient samples, with most lacking temporal information.

This validation demonstrates that our analytical framework successfully distinguishes between different types of genetic variation. The minimal overlap between recombination events and established clusters confirms that cgDist’s enhanced resolution capabilities remain methodologically sound while the integrated recombination detection provides technical capability for comprehensive genomic characterization when required. The cgDist algorithm supports configurable parameters, such as SNP density thresholds (e.g., 3.0%), to accommodate the recombination patterns observed in diverse bacterial pathogens. For example, studies have reported SNP densities ranging from 0.46 to 0.54 SNPs per kilobase in *Clostridioides difficile* strains [Ruppitsch et al., 2015b], and up to 36% of loci exhibiting recombinant variants in soil bacteria [Olm et al., 2020].

### 4.4 Performance Characteristics and Scalability

Comprehensive benchmarking reveals that genetic diversity, assessed through the number of unique allele pairs requiring alignment computation, rather than dataset size, determines computational requirements and optimal threading strategies. High-diversity datasets achieve best performance at 16-32 threads due to memory access contention, while low-diversity datasets maintain near-linear scaling to 128 threads. The Se-1540 *Salmonella* dataset achieved 18.1*×* speedup with 112.9% efficiency at 16 threads, while the Lm-1426 *Listeria* dataset reached 125.1*×* speedup with 97.7% efficiency at 128 threads.

The unified cache architecture provides cumulative benefits for operational surveillance. Progressive cache accumulation in incremental processing validation demonstrated efficiency gains from 0% to 88.3% cache hit rates, with the 300-sample batch processing faster than the 250-sample batch due to alignment reuse. This counter-intuitive scaling behavior, where larger datasets process faster than smaller ones, validates the practical utility of persistent alignment caching for continuous surveillance workflows [Deneke et al., 2021].

### 4.5 Operational Deployment and Multi-Scale Surveillance

The successful integration into the GENPAT bioinformatics platform demonstrates cgDist’s transition from research tool to production infrastructure, with automated Nextflow pipeline achieving sub-minute execution times for cached operations. The production workflow integrates cgDist distance calculations with ReporTree for cluster analysis [Mixão et al., 2023] and SPREAD for interactive cluster visualization [de Ruvo et al., 2024], providing a complete surveillance solution from distance calculation through epidemiological interpretation.

Our validated three-tier surveillance workflow optimizes computational resources while maximizing analytical capability: Tier 1-2 focuses on distance-based clustering for outbreak investigation, while Tier 3 provides recombination analysis for comprehensive population characterization. This graduated approach addresses computational constraints while providing enhanced discrimination where most important for public health decisions.

### 4.6 Community-Shared Cache Infrastructure

A particularly promising direction involves cgDist’s capability to generate comprehensive caches directly from cgMLST schemas without requiring actual isolate data. This enables creation of pre-computed schema-wide caches containing alignments for all possible allele pairs within a schema, which could be hosted on centralized servers and downloaded by laboratories worldwide. Such community-shared caches would eliminate initial computation barriers for new users, as laboratories could simply download pre-computed caches for their cgMLST schemas and immediately begin high-resolution surveillance without the computational overhead of building alignment data from scratch.

This transformation of cache data from institutional assets to community resources would democratize access to nucleotide-level analysis capabilities, particularly benefiting resource-constrained laboratories. The standardized nature of cgMLST schemas ensures cache compatibility across institutions, while the modular cache architecture enables selective downloading of relevant schema subsets, making this approach scalable across diverse surveillance contexts. Community-shared caches could be generated using the most common alignment parameters, ensuring broad compatibility while maintaining the flexibility for specialized applications requiring custom alignment settings.

### 4.7 Comparison with Alternative Approaches and Limitations

While direct performance comparison with traditional SNP-calling pipelines was beyond this study’s scope, cgDist offers conceptual advantages for operational surveillance. Traditional approaches require processing raw sequencing data with substantial computational infrastructure, whereas cgDist leverages pre-computed cgMLST profiles for nucleotide-level resolution without genome reprocessing requirements. The standardized nature of cgMLST loci facilitates inter-laboratory comparability, while cgDist’s incremental processing capability contrasts with SNP-calling’s typical requirement for complete dataset reanalysis [Davis et al., 2015, Gardner et al., 2015].

However, several limitations warrant consideration. cgDist’s resolution remains constrained by cgMLST schema quality and comprehensiveness, though compatibility with wgMLST schemas provides enhanced discrimination for specialized applications. Optimal epidemiological thresholds established for cgMLST distance-based clustering may require recalibration for cgDist analyses, necessitating species-specific validation against known epidemiological relationships [Palma et al., 2022, Ruppitsch et al., 2015a].

Future developments should focus on expanding validation to additional bacterial species to establish species-specific epidemiological thresholds, GPU acceleration for large-scale analyses, and establishing community cache-sharing infrastructure. A particularly promising direction involves cgDist’s capability to generate comprehensive caches directly from cgMLST schemas without requiring actual isolate data. This enables creation of pre-computed schema-wide caches containing alignments for all possible allele pairs within a schema, which could be hosted on centralized servers and downloaded by laboratories worldwide. Such community-shared caches would eliminate initial computation barriers for new users, as laboratories could simply download pre-computed caches for their cgMLST schemas and immediately begin high-resolution surveillance without the computational overhead of building alignment data from scratch. This transformation of cache data from institutional assets to community resources would democratize access to nucleotide-level analysis capabilities, particularly benefiting resource-constrained laboratories. Extension to amino acid sequence analysis would further enable functional context for genetic variations, particularly valuable for antimicrobial resistance and virulence factor surveillance.

### 4.8 Public Health Impact and Future Directions

The prevention of false source attribution demonstrated through validated priority reorderings has immediate implications for public health policy and regulatory enforcement. Enhanced discrimination reduces risks of incorrectly implicating food production facilities or environmental sources, potentially preventing unnecessary facility closures, product recalls, and regulatory actions with substantial economic consequences [Maia et al., 2021]. The ability to resolve tied cgMLST distances into distinct priority levels enables evidence-based resource allocation, maximizing probability of identifying true transmission pathways early in outbreak investigations.

cgDist contributes to genomic epidemiology’s evolution toward real-time surveillance by bridging traditional and advanced methodologies without requiring complete system reconstruction. The standardized output formats and compatibility with established pipelines enable gradual integration while preserving institutional knowledge and analytical continuity [Didelot et al., 2017].

As bacterial surveillance systems continue advancing toward sophisticated analytical capabilities, cgDist provides both immediate practical benefits and a methodological framework for further development. The demonstrated species-independent benefits, preserved epidemiological interpretation frameworks, and evolutionary rather than revolutionary implementation approach facilitate institutional adoption while delivering enhanced analytical capabilities that directly benefit public health outcomes.

### 4.9 Conclusions

cgDist successfully addresses the fundamental tension between computational efficiency and genetic resolution in bacterial genomic surveillance, providing nucleotide-level discrimination within the operational constraints of routine public health practice. The cluster zoom paradigm establishes cgDist’s optimal use case: serving as a precision “zoom lens” for the detailed investigation of epidemiologically relevant clusters identified through initial cgMLST screening. This targeted resolution enhancement strategy maximizes epidemiological insight precisely where it matters most for outbreak investigation, rather than attempting wholesale restructuring of global population relationships.

The algorithm’s unified cache architecture transforms genomic surveillance from batch-based processing to continuous streaming analysis, with cumulative performance benefits that improve operational efficiency over time. The prevention of false source attribution through enhanced discrimination has immediate implications for regulatory enforcement and public health policy, reducing risks of unnecessary facility closures and product recalls while enabling evidence-based resource allocation during outbreak investigations.

The successful integration into operational surveillance systems validates cgDist’s transition from research tool to production infrastructure, with automated workflows achieving sub-minute execution times and real-time outbreak detection capabilities. The species-independent benefits, preserved epidemiological frameworks, and gradual integration approach facilitate institutional adoption while providing a methodological foundation for continued innovation in computational genomic epidemiology.

cgDist represents a significant methodological advancement that bridges the resolution gap between traditional cgMLST and computationally intensive approaches, delivering enhanced analytical capabilities that directly benefit public health outcomes through improved accuracy in outbreak investigation and source attribution.

## Data Availability

All datasets used in this study are publicly available through the BeONE project (Bacterial pathogens in the One Health European Joint Programme) repositories. The *Listeria monocytogenes* datasets (Lm-1426 and Lm-1874) and *Salmonella enterica* datasets (Se-1434 and Se-1540) can be accessed through Zenodo at https://doi.org/10.5281/zenodo.7120166 and https://doi.org/10.5281/zenodo.7267822. Raw sequencing data and assembled genomes are available through the European Nucleotide Archive (ENA) under the accession numbers provided in the BeONE metadata files. The cgMLST schemas used for analysis are the INNUENDO schemas [Moura et al., 2017], available through ChewieNS (https://chewbbaca.online/) for *Listeria monocytogenes* (accessed March 2024) and *Salmonella enterica* (accessed March 2024). For *Salmonella enterica*, we applied the EFSA-filtered subset (3,255 loci) following European food safety surveillance standards, available from https://doi.org/10.5281/zenodo.1323684. Cache files generated during this study are available upon request from the corresponding author due to their large file sizes. The star clustering analysis results and epidemiological metadata used for source prioritization analysis are provided as supplementary datasets.

All supplementary data files, including distance matrices (Hamming, cgDist SNPs-only, SNPs+InDel-events, and SNPs+InDel-bases modes), allelic profiles, and recombination analysis results, are publicly available on Zenodo at https://doi.org/10.5281/zenodo.17285518. Validation scripts are available upon request from the corresponding author.

## Code Availability

The cgDist software is open-source and freely available under the MIT license. The core algorithm implementation in Rust is available at https://github.com/genpat-it/cgDist, with comprehensive documentation including installation instructions, usage examples, and API reference. A Docker image for containerized deployment will be made publicly available upon publication at Docker Hub (cgdist/cgdist:latest).

## 5 Author Contributions

Andrea de Ruvo conceived and led the development of the `cgDist` algorithm, served as the primary author, and acted as the corresponding author for the study. Michele Flammini provided PhD supervision, theoretical guidance, and methodological oversight throughout the research. Pierluigi Castelli managed the preparation and curation of the datasets used in the analysis, ensuring their quality and accessibility. Andrea Bucciacchio, as the DevOps engineer, designed and maintained the platform infrastructure that supported the execution of the genomic pipelines. Iolanda Mangone and Nicolas Radomski, as bioinformatics experts, provided critical supervision and guidance throughout the algorithm’s development, ensuring its scientific robustness. Verónica Mixão and Vítor Borges provided valuable suggestions that significantly improved the `cgDist` algorithm, enhancing its functionality and applicability. Adriano Di Pasquale, head of the bioinformatics unit at the Istituto Zooprofilattico Sperimentale dell’Abruzzo e del Molise (Teramo), facilitated the research by providing essential resources and institutional support. All authors contributed to the writing and revision of the manuscript and approved the final version.

## 6 Acknowledgments

We express our gratitude to the Gran Sasso Science Institute (GSSI), L’Aquila, where Andrea de Ruvo is pursuing his PhD, for providing academic support and resources. We sincerely thank the Istituto Zooprofilattico Sperimentale dell’Abruzzo e del Molise, Teramo, for supplying essential resources that made this research possible. We also acknowledge the Instituto Nacional de Saúde Doutor Ricardo Jorge (INSA), Lisbon, for their valuable suggestions and feedback, which significantly contributed to the development of the `cgDist` algorithm.

## Supplementary Information

This supplementary information document contains three main sections: data availability, detailed methodological descriptions, and additional results. All materials are organized for clarity and ease of access.

## Supplementary Data

The following supplementary data files are available with this article, organized by species for clarity and ease of access:

## Listeria monocytogenes Dataset 1426

**File:** `Lm1_426_allelic_profiles.tsv` - Allelic profile matrix (1,426 samples). Contains cgMLST profiles for all samples used in the analysis. File format: TSV.

**File:** `Lm1426_hamming.tsv` - Hamming distance matrix (1,426 samples). Contains pairwise Hamming distances calculated from allelic profiles using cgmlst-dists. File format: TSV.

**File:** `Lm1426_hamming_98percent.tsv` - Hamming distance matrix (98% sample completeness threshold). Filtered dataset containing only samples with at least 98% of loci successfully called. File format: TSV.

**File:** `Lm1426_cgdist _snps _indel_events.tsv` - cgDist matrix - SNPs + InDel events (1,426 samples). Contains pairwise distances calculated using the cgDist algorithm counting SNPs and InDel events. File format: TSV.

**File:** `Lm1426_cgdist_snps _indel_events _98percent.tsv` - cgDist matrix - SNPs + InDel events (98% sample completeness threshold). Filtered dataset containing only samples with at least 98% of loci successfully called. File format: TSV.

**File:** `Lm1426_cgdist_snps_only.tsv` - cgDist matrix - SNPs only (1,426 samples). Contains pairwise distances calculated using only SNPs. File format: TSV.

**File:** `Lm1426 _cgdist_snps _only_98percent.tsv` - cgDist matrix - SNPs only (98% sample completeness threshold). Filtered dataset containing only samples with at least 98% of loci successfully called. File format: TSV.

**File:** `Lm1426_cgdist _snps _indel _bases.tsv` - cgDist matrix - SNPs + InDel bases (1,426 samples). Contains pairwise distances calculated using SNPs and total InDel bases. File format: TSV.

**File:** `Lm1426 _cgdist_snps _indel_bases _98percent.tsv` - cgDist matrix - SNPs + InDel bases (98% sample completeness threshold). Filtered dataset containing only samples with at least 98% of loci successfully called. File format: TSV.

**File:** `Lm1426_recombination_analysis.tsv` - Legacy recombination analysis results. File format: TSV.

**File:** `Lm1426 _recombination analysis_with _samples_98percent.tsv` - Detailed recom-bination events analysis demonstrating strict clonal inheritance patterns in *Listeria monocytogenes* populations (98% completeness threshold). Contains individual recombination events with complete metrics, sample pair identifiers, locus information, SNP/InDel counts, sequence lengths, and mutation density calculations. File format: TSV.

**File:** `Lm1426 _pairwise _recombination_summary_98percent.tsv` - Pairwise recombination summary for *Listeria monocytogenes* dataset (98% completeness threshold). Provides sample pairs with total recombining loci counts, percentage of analyzed loci showing recombination signatures, and summary statistics for epidemiological interpretation. File format: TSV.

## Listeria monocytogenes Dataset 1874

**File:** `Lm1874_allelic _profiles.tsv` - Allelic profile matrix (1,874 samples). Contains cgMLST profiles for the expanded dataset. File format: TSV.

**File:** `Lm1874_hamming.tsv` - Hamming distance matrix (1,874 samples). File format: TSV.

**File:** `Lm1874_hamming _98percent.tsv` - Hamming distance matrix (98% sample completeness threshold). Filtered dataset containing only samples with at least 98% of loci successfully called. File format: TSV.

**File:** `Lm1874_cgdist_snps_indel _events.tsv` - cgDist matrix - SNPs + InDel events (1,874 samples). File format: TSV.

**File:** `Lm1874_cgdist _snps_indel _events _98percent.tsv` - cgDist matrix - SNPs + InDel events (98% sample completeness threshold). Filtered dataset containing only samples with at least 98% of loci successfully called. File format: TSV.

**File:** `Lm1874 _cgdist_snps _only.tsv` - cgDist matrix - SNPs only (1,874 samples). File format: TSV.

**File:** `Lm1874_cgdist_snps _only_98percent.tsv` - cgDist matrix - SNPs only (98% sample completeness threshold). Filtered dataset containing only samples with at least 98% of loci successfully called. File format: TSV.

**File:** `Lm1874_cgdist _snps _indel_bases.tsv` cgDist matrix - SNPs + InDel bases (1,874 samples). File format: TSV.

**File:** `Lm1874 _cgdist_snps_indel_bases _98percent.tsv` - cgDist matrix - SNPs + InDel bases (98% sample completeness threshold). Filtered dataset containing only samples with at least 98% of loci successfully called. File format: TSV.

**File:** `Lm1874 _recombination_analysis _with_samples _98percent.tsv` - Detailed recombination events analysis demonstrating strict clonal inheritance patterns in expanded *Listeria monocytogenes* populations (98% completeness threshold). Contains individual recombination events with complete metrics, sample pair identifiers, locus information, SNP/InDel counts, sequence lengths, and mutation density calculations. File format: TSV.

**File:** `Lm1874 _pairwise _recombination _summary_98percent.tsv` - Pairwise recombination summary for expanded *Listeria monocytogenes* dataset (98% completeness threshold). Provides sample pairs with total recombining loci counts, percentage of analyzed loci showing recombination signatures, and summary statistics for epidemiological interpretation. File format: TSV.

## Salmonella enterica Dataset 1540

**File:** `Se1540_allelic_profiles.tsv` - Allelic profile matrix (1,540 samples). Contains cgMLST profiles for all samples used in the analysis. File format: TSV.

**File:** `Se1540_hamming.tsv` - Hamming distance matrix (1,540 samples). Contains pairwise Hamming distances calculated from allelic profiles. File format: TSV.

**File:** `Se1540_hamming _98percent.tsv` - Hamming distance matrix (98% sample completeness threshold). Filtered dataset containing only samples with at least 98% of loci successfully called. File format: TSV.

**File:** `Se1540_cgdist_snps_indel _events.tsv` cgDist matrix - SNPs + InDel events (1,540 samples). Contains pairwise distances calculated using the cgDist algorithm. File format: TSV.

**File:** `Se1540_cgdist_snps_indel _events _98percent.tsv` - cgDist matrix - SNPs + InDel events (98% sample completeness threshold). Filtered dataset containing only samples with at least 98% of loci successfully called. File format: TSV.

**File:** `Se1540 _cgdist_snps _only.tsv` - cgDist matrix - SNPs only (1,540 samples). Contains pairwise distances calculated using only SNPs. File format: TSV.

**File:** `Se1540 _cgdist _snps_only_98percent.tsv` - cgDist matrix - SNPs only (98% sample completeness threshold). Filtered dataset containing only samples with at least 98% of loci successfully called. File format: TSV.

**File:** `Se1540 _cgdist_snps_indel_bases.tsv` - cgDist matrix - SNPs + InDel bases (1,540 samples). Contains pairwise distances calculated using SNPs and total InDel bases. File format: TSV.

**File:** `Se1540_cgdist _snps _indel_bases_98percent.tsv` - cgDist matrix - SNPs + InDel bases (98% sample completeness threshold). Filtered dataset containing only samples with at least 98% of loci successfully called. File format: TSV.

**File:** `Se1540_recombination_analysis.tsv` - Legacy recombination analysis results. File format: TSV.

**File:** `Se1540_recombination_analysis_with _samples_98percent.tsv` - Detailed recombination events analysis for *Salmonella enterica* dataset (98% completeness threshold). Contains individual recombination events affecting 20 sample pairs (0.03% of EFSA loci) with complete metrics, sample pair identifiers, locus information, SNP/InDel counts, sequence lengths, and mutation density calculations detected using 3.0% threshold. File format: TSV.

**File:** `Se1540_pairwise _recombination_summary_98percent.tsv` - Pairwise recombination summary for *Salmonella enterica* dataset (98% completeness threshold). Provides sample pairs with total recombining loci counts, percentage of analyzed loci showing recombination signatures, and summary statistics for epidemiological interpretation. File format: TSV.

**File:** `Se1540_mutation_density _analysis.png` - Comprehensive visualization of mutation density distribution analysis supporting the empirical validation of the 3.0% recombination threshold. Includes histogram, cumulative distribution, threshold ranges analysis, and log-scale distribution plots. File format: PNG.

## Salmonella enterica Dataset 1434

**File:** `Se1434 _allelic _profiles.tsv` - Allelic profile matrix (1,434 samples). Contains cgMLST profiles for the validation dataset. File format: TSV.

**File:** `Se1434_hamming.tsv` - Hamming distance matrix (1,434 samples). File format: TSV.

**File:** `Se1434 _hamming_98percent.tsv` - Hamming distance matrix (98% sample completeness threshold). Filtered dataset containing only samples with at least 98% of loci successfully called. File format: TSV.

**File:** `Se1434_cgdist_snps _indel _events.tsv` - cgDist matrix - SNPs + InDel events (1,434 samples). File format: TSV.

**File:** `Se1434 _cgdist_snps _indel _events_98percent.tsv` - cgDist matrix - SNPs + InDel events (98% sample completeness threshold). Filtered dataset containing only samples with at least 98% of loci successfully called. File format: TSV.

**File:** `Se1434_cgdist _snps_only.tsv` - cgDist matrix - SNPs only (1,434 samples). File format: TSV.

**File:** `Se1434_cgdist_snps_only _98percent.tsv` - cgDist matrix - SNPs only (98% sample completeness threshold). Filtered dataset containing only samples with at least 98% of loci successfully called. File format: TSV.

**File:** `Se1434 _cgdist _snps_indel _bases.tsv` - cgDist matrix - SNPs + InDel bases (1,434 samples). File format: TSV.

**File:** `Se1434 _cgdist _snps _indel _bases _98percent.tsv` - cgDist matrix - SNPs + InDel bases (98% sample completeness threshold). Filtered dataset containing only samples with at least 98% of loci successfully called. File format: TSV.

**File:** `Se1434 _recombination_analysis.tsv` - Legacy recombination analysis results for validation dataset. File format: TSV.

**File:** `Se1434 _recombination _analysis_with _samples_98percent.tsv` - Detailed recombination events analysis for validation *Salmonella enterica* dataset (98% completeness threshold). Contains minimal recombination activity (1 sample pair) with complete metrics, sample pair identifiers, locus information, SNP/InDel counts, sequence lengths, and mutation density calculations detected using 3.0% threshold. File format: TSV.

**File:** `Se1434_pairwise _recombination _summary _98percent.tsv` - Pairwise recombination summary for validation *Salmonella enterica* dataset (98% completeness threshold). Provides sample pairs with total recombining loci counts, percentage of analyzed loci showing recombination signatures, and summary statistics for epidemiological interpretation. File format: TSV.

## Summary

All supplementary data files are provided in Tab-Separated Values (TSV) format for maximum compatibility with computational biology software. Distance matrices include metadata headers with analysis parameters and timestamps for reproducibility. The complete dataset enables researchers to reproduce all analyses presented in this study and conduct their own comparative studies.

Files are organized by species and dataset size with consistent naming conventions. The allelic profiles provide the raw cgMLST data serving as input for all distance calculations, while Hamming matrices contain traditional allelic difference calculations establishing baseline comparisons. The cgDist matrices encompass three calculation modes: SNPs+InDel events counting both SNPs and individual InDel events, SNPs only focusing exclusively on single nucleotide changes, and SNPs+InDel bases weighting InDels by total affected nucleotides. Files designated with the 98% threshold suffix represent high-quality subsets filtered for samples with at least 98% of loci successfully called. The Se-1540 dataset uniquely includes recombination analysis results demonstrating cgDist’s horizontal gene transfer detection capabilities, accompanied by mutation density analysis providing statistical validation of empirically-derived recombination thresholds.

The comprehensive dataset supports reproducible research and enables the scientific community to evaluate cgDist performance across multiple pathogen species and analytical scenarios. Each species section contains complete analytical workflows from raw allelic profiles through all distance calculation modes, enabling researchers to reproduce analyses or adapt methods to their own surveillance datasets. Additionally, the Se-1540 dataset includes recombination analysis results demonstrating cgDist’s integrated horizontal gene transfer detection capabilities, providing both detailed event data and empirical threshold validation for advanced genomic surveillance applications.

## Supplementary Methods

### Debugging and Quality Assurance Capabilities

While cgDist’s standard operation focuses on efficient aggregate distance metrics, the tool architecture supports detailed debugging and quality assurance through optional preservation of alignment details. Users can optionally enable verbose logging and detailed alignment output that captures the complete Parasail alignment results, including alignment matrices, trace-back paths, and variant positions. This debugging capability enables researchers to manually inspect the specific nucleotide differences contributing to distance calculations, validate alignment quality, and identify potential sequencing or assembly artifacts that might affect epidemiological interpretation.

The debugging features encompass complete alignment matrices showing the dynamic programming calculations, trace-back paths indicating the best alignment route through the scoring matrix, detailed variant positions with nucleotide-level annotations for each detected difference, per-locus alignment statistics and quality metrics enabling identification of problematic regions, and comprehensive validation reports for cache integrity and consistency verification.

However, enabling comprehensive alignment debugging significantly increases storage requirements and computational overhead, making it suitable primarily for validation studies, method development, or troubleshooting specific analytical scenarios rather than routine surveillance applications. For routine surveillance, the standard mode provides sufficient information through summary statistics while maintaining good performance.

### Detailed Distance Calculation Algorithm

The following algorithm provides the complete technical details for analyzing pairwise sequence alignments and computing different distance metrics from the classified variants.

#### Algorithm 2: Distance Calculation from Sequence Alignment

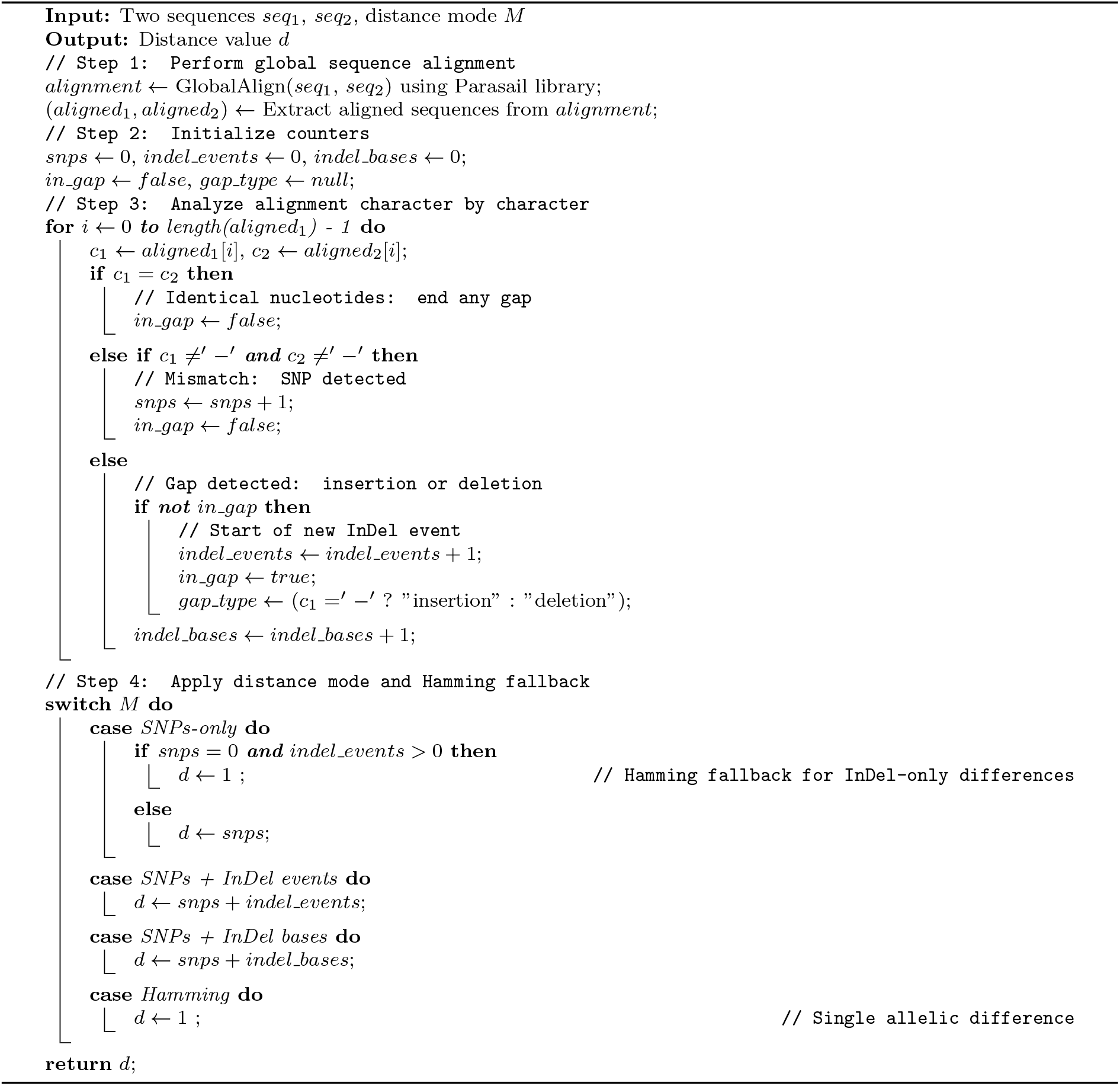

### Alignment Parameters and Technical Details

#### Alignment Parameter Settings

cgDist employs the DNA-strict alignment mode with a match score of +3, mismatch penalty of −2, gap opening penalty of 8, and gap extension penalty of 3.

This predefined parameter set is optimized for bacterial genomic sequences, with elevated gap penalties designed to favor SNP-based explanations over complex InDel patterns while minimizing alignment artifacts. The conservative parameter selection enhances discriminatory power at epidemiologically important distance ranges where outbreak investigation decisions are most sensitive.

#### Implementation Details

The cgDist algorithm is implemented in Rust 2021 edition for memory safety and performance, utilizing Parasail-rs v0.3.5 Rust bindings for sequence alignment with SIMD vectorization support. The system employs native Rust threading with a work-stealing scheduler for parallelization, LZ4 algorithm for efficient cache compression, and compressed JSON format with metadata headers for cache storage. Supported hashers include CRC32 (default), MD5, SHA256, sequence-based, and Hamming algorithms.

The Parasail library provides optimized C implementation with Rust bindings, ensuring both computational efficiency and memory safety while maintaining deterministic alignment results required for reproducible distance calculations.

#### Detailed Computational Complexity Analysis

To analyze cgDist computational requirements, we perform formal complexity analysis examining time complexity *T* (*n, s, l*) and space complexity *S*(*n, u, l*), where *n* represents the number of genomic samples, *s* the maximum nucleotide sequence length per locus in the schema, *l* the number of cgMLST loci, and *u* the average number of unique alleles per locus in the schema.

### Time Complexity Analysis

#### Upper Bound

The algorithm computes pairwise distances for 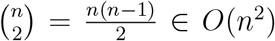 unique sample pairs. For each pair (*i, j*), we compare *l* cgMLST loci, performing sequence alignment when alleles differ. The alignment cost per locus is *O*(*s*^2^) for the worst-case maximum sequence length using the Needleman-Wunsch algorithm. Therefore:

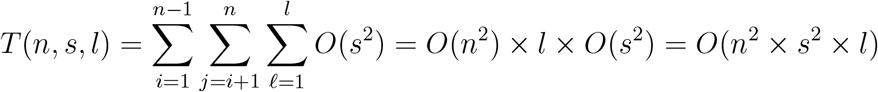

#### Lower Bounds

The algorithm must examine each sample pair at least once (Ω(*n*^2^)) and process all loci for meaningful distances (Ω(*l*)), establishing *T* (*n, s, l*) ∈ Ω(*n*^2^ *× l*). In the worst case where all alleles differ, sequence alignment adds Ω(*s*^2^) per comparison, yielding the tight bound *T* (*n, s, l*) ∈ Θ(*n*^2^ *× l × s*^2^).

#### Optimization

Cache optimization reduces effective time complexity to *O*(*n*^2^ *× l*) with high hit rates, transforming *O*(*s*^2^) alignment operations into *O*(1) lookups. Parallelization achieves *T*_parallel_ = *O*(*n*^2^ *× s*^2^ *× l/p*) for *p* processors, limited by Amdahl’s law and cache contention.

### Space Complexity Analysis

#### Memory Requirements

Memory requirements include the symmetric distance matrix *O*(*n*^2^), alignment cache storage *O*(*l × u*^2^) for allele pair alignments, allelic profiles *O*(*n × l*), and temporary alignment buffers *O*(*s*^2^) for dynamic programming matrices (allocated per alignment operation). The dominant term depends on dataset characteristics:

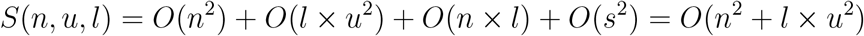

#### Dominance Cases

For small datasets with high allelic diversity (*n*^2^ *< l × u*^2^), cache storage dominates memory usage. For large sample collections with moderate diversity (*n*^2^ *> l × u*^2^), the distance matrix becomes the primary memory constraint.

#### Practical Performance Considerations

The actual performance of cgDist depends heavily on several factors including cache hit rates, genetic diversity, sequence length distribution, and parallelization efficiency. In bacterial surveillance with clonal population structures, many allele pairs are shared across samples, leading to high cache utilization. Highly diverse populations require more unique alignments but benefit more from resolution enhancement, while shorter sequences reduce alignment costs while maintaining epidemiological utility. Parallelization efficiency depends on work distribution across processors with minimal cache contention.

### Integrated Recombination Analysis Details

#### Recombination Detection Framework

cgDist’s unified cache architecture naturally enables integrated recombination analysis by leveraging alignment statistics already computed during standard distance calculations. When cgDist caches are enriched with nucleotide sequence length information from the schema using the `--enrich-lengths` parameter, the system can identify potential recombination events through mutation density analysis without additional computational overhead. This enrichment process stores the original sequence lengths alongside alignment statistics in the cache, enabling mutation density calculations 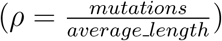 for recombination detection in subsequent analyses.

The recombination detection algorithm operates on the principle that genetic recombination events typically exhibit elevated mutation densities compared to vertical transmission, where mutations accumulate gradually over time through point mutations and small indels. By analyzing the ratio of total mutations (SNPs + InDel events) to average sequence length between allele pairs, the method can identify loci that may have undergone horizontal gene transfer or other recombination processes.

#### Algorithmic Implementation

The recombination analysis utilizes three key filtering steps: epidemiological filtering, mutation density analysis, and loci filtering. Epidemiological filtering restricts analysis to sample pairs within specified Hamming distance thresholds (≤ 15 alleles, empirically determined from dataset analysis) to focus on epidemiologically relevant comparisons that could represent recent transmission or clonal expansion. Mutation density analysis calculates mutation density as 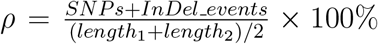 for each allele pair, where sequences exceeding the threshold (3.0%) are flagged as potential recombination events. Loci filtering applies optional loci subsets (e.g., EFSA core loci for *Salmonella enterica*) to focus analysis on epidemiologically relevant genomic regions.

#### Algorithm Workflow

The algorithm processes enriched cache entries through a systematic workflow that begins by loading enriched cache containing alignment statistics and sequence lengths, then parsing allelic profiles to map CRC hashes to samples. The system calculates Hamming distances between sample pairs and filters pairs by epidemiological relevance using the Hamming threshold. For each qualifying pair, the algorithm analyzes all shared loci by calculating mutation density for each allele pair, comparing against the recombination threshold, and recording events exceeding the threshold. Finally, the system generates detailed event reports and pairwise summaries for epidemiological interpretation.

#### Output Files and Interpretation

The recombination analysis generates two complementary output files:

#### Detailed Events File

({`Dataset _recombination_analysis_with_samples _98percent.tsv`}) contains individual recombination events with complete metrics for samples meeting 98% completeness threshold. Includes sample pair identifiers and locus information, SNP counts, InDel counts, sequence lengths, and density calculations, along with allele CRC identifiers for traceability.

#### Pairwise Summary File

({`Dataset_pairwise_recombination_summary_98percent.tsv`}) provides sample pairs with total recombining loci counts for the 98% completeness subset, percentage of analyzed loci showing recombination signatures, and summary statistics for epidemiological interpretation.

#### Practical Implementation

Cache enrichment is performed during the initial analysis by adding the `--enrich-lengths` flag to standard cgDist execution. Once enriched, the cache contains both alignment statistics and sequence length data, enabling recombination analysis in subsequent runs using the `--recombination-log` parameter without additional computational overhead.

This design allows laboratories to enable recombination detection capabilities retroactively on existing analyses by enriching their established cache files. The modular approach ensures that recombination analysis can be added to existing workflows without disrupting established analytical pipelines.

#### Scientific Applications

The integrated recombination analysis enables several important scientific applications including outbreak investigation to detect recombination events within transmission chains that could affect phylogenetic inference, food safety surveillance to monitor horizontal gene transfer in pathogen populations, evolutionary analysis to track population structure changes and gene flow patterns, antimicrobial resistance monitoring to detect transfer of resistance genes between bacterial lineages, and epidemiological clustering to identify samples that may require specialized phylogenetic methods due to recombination.

#### Recombination Analysis Algorithm

The following pseudocode describes the complete recombination detection algorithm implemented in cgDist:

##### Algorithm 3: Recombination Analysis from Enriched Cache

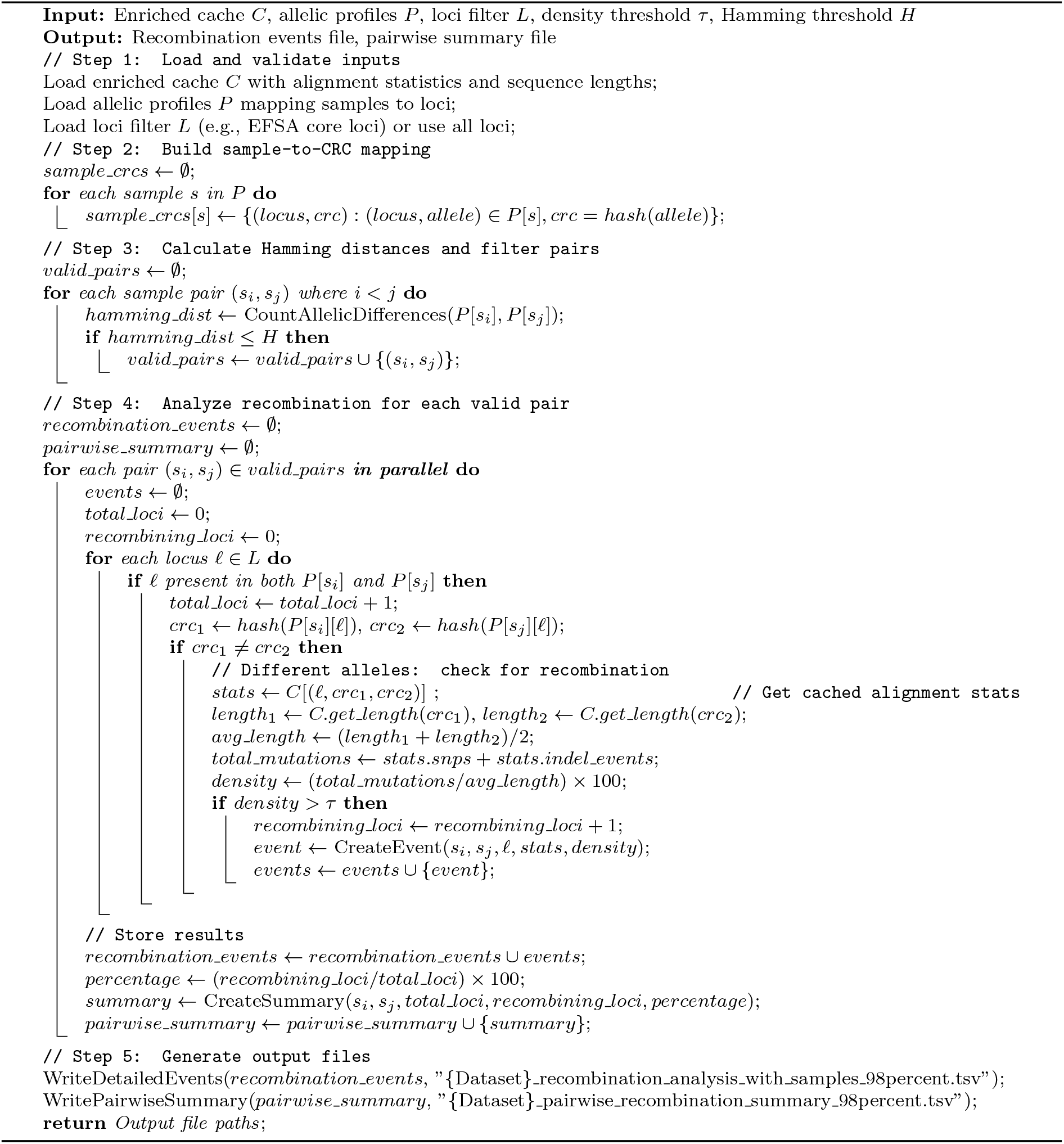

## Supplementary Results

This section will contain additional results, extended analyses, and detailed performance comparisons that support the main findings presented in the manuscript.

### Scalability and Performance Analysis

We conducted comprehensive scalability analysis using high-performance computing infrastructure to evaluate cgDist performance characteristics across different dataset sizes and genetic diversity levels. Testing was performed on a 128-core server with 720 GB RAM running AlmaLinux 8.10, including thread scaling from 1 to 128 cores, cache performance measurement across multiple runs, and comparison with existing tools.

### Parallelization Performance and Dataset-Dependent Scalability Patterns

cgDist demonstrates dataset-dependent scalability patterns that correlate strongly with genetic diversity rather than dataset size. Comprehensive scalability testing revealed two distinct performance profiles reflecting the underlying biological characteristics of the analyzed populations.

For the high-diversity Se-1540 *Salmonella enterica* dataset (6.05M unique pairs, genetic diversity index 0.037), best performance occurs at 16-32 threads with diminishing returns beyond this range. The best configuration at 16 threads delivers 18.1 *×* speedup with 112.9% efficiency, achieving peak throughput of 2,077 pairs/second. While maximum throughput of 3,939 pairs/second is achieved at 64 threads, efficiency drops substantially to 53.5%. Beyond 64 threads, severe thread contention results in performance collapse, with 128 threads showing only 0.4% efficiency. Memory requirements remain stable at 3.5 GB across all thread configurations, indicating that memory usage is independent of parallelization strategy.

In contrast, the low-diversity Lm-1426 *Listeria monocytogenes* dataset (0.72M unique pairs, genetic diversity index 0.018) exhibits near-linear scaling across the entire thread range. The system achieves 125.1*×* speedup at 128 threads with 97.7% efficiency, maintaining consistent per-thread performance of approximately 42 pairs/second across all configurations. Peak performance reaches 5,378 pairs/second at maximum thread utilization without performance degradation. Memory efficiency is notable, requiring only 1.9 GB (45% reduction compared to high-diversity datasets), demonstrating that genetic diversity rather than dataset size determines memory requirements.

### Unified Distance Cache Architecture Performance

cgDist’s unified distance cache system provides substantial performance improvements for multi-mode analysis workflows, particularly important for surveillance applications requiring multiple distance calculation modes. Cache performance evaluation across both species using production datasets demonstrates the significant efficiency gains achievable through intelligent caching strategies.

The cache architecture exhibits notable performance differentials between cache-building and cache-utilization phases. For the *Salmonella enterica* Se-1540 dataset (6.05M pairs), initial SNP analysis with cache building processes 2,007 pairs/second requiring 50:15 execution time and 5.57 GB memory. However, subsequent SNPs+indel-events analysis using cached alignments achieves 28,374 pairs/second (14*×* improvement) with significantly reduced execution time of 3:33 and 29% memory reduction to 3.95 GB. Similarly, SNPs+indel-bases analysis maintains 20,938 pairs/second processing rate with comparable memory efficiency.

For the *Listeria monocytogenes* Lm-1874 dataset (4.26M pairs), cached SNP analysis achieves 25,900 pairs/second completing in 2:44 with 3.18 GB memory usage. When building cache for SNPs+indel-events analysis, processing rate drops to 1,617 pairs/second requiring 43:54 execution time and 4.51 GB memory. Subsequent cached SNPs+indel-bases analysis recovers to 20,470 pairs/second with reduced memory requirements, demonstrating 94% time reduction when reusing caches across analysis modes.

### Threading Performance Analysis

Performance scaling was evaluated across thread counts from 1 to 128 cores using representative datasets from each species. The analysis measured alignment throughput as unique allele pairs processed per second, memory utilization including peak and average memory consumption patterns, cache efficiency through hit rates and lookup performance across thread counts, and load balancing via work distribution across available processors.

### Comparative Performance Benchmarking

cgDist performance was compared against existing tools including cgmlst-dists for baseline cgMLST analysis and representative SNP-calling pipelines. Benchmarks included computation time as wall-clock time for distance matrix generation, memory requirements measured as peak memory usage across dataset sizes, accuracy validation through consistency with established distance metrics, and scalability limits determining maximum practical dataset sizes.

### Incremental Processing Validation

To demonstrate practical deployment capability for real-time surveillance, we developed a comprehensive Nextflow pipeline for incremental processing validation. This pipeline simulates the temporal arrival of isolates in surveillance scenarios and measures the computational performance, cache efficiency, and cumulative benefits of alignment reuse as surveillance programs continuously integrate new genomic data.

### Nextflow Pipeline Architecture

The incremental processing pipeline implements a systematic workflow that establishes initial cache by processing the first dataset to build baseline cache, performs incremental updates by sequentially adding new isolates simulating real-time arrival, conducts performance monitoring to track computation time, memory usage, and cache statistics, executes validation checks to ensure distance matrix consistency across incremental updates, and measures efficiency metrics including cache hit rates and alignment reuse benefits.

### Experimental Design

The validation approach enables assessment of incremental processing performance and cache efficiency benefits in realistic surveillance deployment scenarios. The pipeline pro-cesses datasets in temporal simulation mode, measuring cache growth patterns to determine how cache size and hit rates evolve with dataset growth, computational efficiency through reduction in processing time for new isolates, memory scaling as surveillance datasets expand, and update latency representing the time required to integrate new isolates into existing analyses.

### Production Surveillance Integration Results

Incremental processing validation confirms that cgDist’s cache-aware design enables practical deployment of nucleotide-level genomic surveillance at the scale and speed required for modern public health practice. Cache hit rates improve progressively from initial runs to subsequent analyses as surveillance programs accumulate alignment data over time, demonstrating the cumulative performance benefits needed for continuous pathogen monitoring workflows.

The practical utility of cgDist extends beyond algorithmic improvements to encompass comprehensive integration into real-world genomic surveillance workflows. cgDist has been successfully integrated into operational surveillance systems and provides a foundation for continuous pathogen monitoring in public health laboratories.

cgDist has been successfully integrated into the GENPAT bioinformatics platform, where it serves as the core distance calculation engine for real-time epidemiological analysis. The integration demonstrates production-ready capabilities including sub-minute execution times for cached operations, efficient incremental updates as new samples are added to surveillance databases, and 94% time reduction when reusing caches across different distance calculation modes.

### Surveillance Performance Validation Results

Comprehensive validation of the cgDist surveillance pipeline using incremental *Salmonella enterica* batches demonstrates cache performance improvements for real-time genomic surveillance deployment (Table 6). The test configuration utilized the cgdist-resources schema with EFSA loci filtering (3,255 loci) in SNPs+InDel-events mode with DNA-strict alignment parameters.

**Table 6:** Progressive cache performance improvements in cgDist surveillance pipeline validation. Incremental batches of *Salmonella enterica* samples demonstrate cache efficiency gains from 0% to 88.3% over six surveillance updates, with the 300-sample batch processing faster than the 250-sample batch due to improved cache utilization.

| Batch | Samples | Time (s) | Cache Hit (%) | Efficiency (%) | Cache Entries | Cache Size |
| --- | --- | --- | --- | --- | --- | --- |
| 1 | 50 | 59.8 | 0.0 | 0.0 | 267,231 | 8.9 MB |
| 2 | 100 | 56.6 | 56.5 | 56.5 | 472,616 | 15.7 MB |
| 3 | 150 | 38.4 | 79.3 | 79.3 | 596,215 | 19.8 MB |
| 4 | 200 | 82.2 | 66.4 | 66.4 | 897,546 | 29.9 MB |
| 5 | 250 | 108.9 | 68.6 | 68.6 | 1,308,063 | 43.6 MB |
| 6 | 300 | 61.3 | 88.3 | 88.3 | 1,481,540 | 49.4 MB |

**Note:** Se-1540 dataset with SNPs+InDel-events mode, DNA-strict alignment parameters, and EFSA schema. Each batch contains cumulative samples (batch 2 includes batch 1 samples plus 50 new). Cache efficiency represents the percentage of alignments reused from previous computations. The counter-intuitive performance improvement at batch 6 (300 samples faster than 250 samples) demonstrates the significant impact of high cache hit rates on surveillance scalability.

Progressive cache accumulation reveals efficiency gains: initial processing of 50 samples required 59.8 seconds with 0% cache efficiency, establishing the baseline cache of 267,231 alignments. The first incremental update to 100 samples achieved immediate performance improvement with 56.5% cache hit rate, reducing processing time to 56.6 seconds despite doubling the sample size. Subsequent batches demonstrated accelerating efficiency gains: 150 samples processed in 38.4 seconds (79.3% cache hit), 200 samples in 82.2 seconds (66.4% cache hit), and 250 samples in 108.9 seconds (68.6% cache hit).

The 300-sample batch completed in 61.3 seconds despite being the largest dataset—faster than the 250-sample batch—due to achieving 88.3% cache hit rate. This counter-intuitive result, where larger datasets process faster than smaller ones, demonstrates the impact of cache-aware architecture on surveillance scalability. The cache grew progressively from 8.9MB (267K alignments) to 49.4MB (1.48M alignments), maintaining compression efficiency through LZ4 encoding with approximately 33 bytes per alignment entry, providing a reusable computational resource for ongoing surveillance operations.

### Integrated Recombination Analysis Results

#### Recombination Detection in *Salmonella enterica* Surveillance

To demonstrate the practical utility of cgDist’s integrated recombination analysis capabilities, we conducted a comprehensive analysis using the *Salmonella enterica* Se-1540 dataset with enriched cache data containing sequence length information. This proof-of-concept analysis demonstrates how cgDist’s unified architecture enables multi-scale genomic surveillance by leveraging existing alignment statistics without additional computational overhead.

#### Empirical Threshold Validation

Comprehensive distribution analysis of Se-1540 recombination events validated the 3.0% mutation density threshold through statistical characterization. The mutation densities exhibited a bimodal distribution (modes at 4.9% and 16.5%) with median density 3.50% and first quartile 2.90%. The 3.0% threshold was positioned near the median and closely aligned with natural population structure, effectively discriminating between typical allelic variation and genuine horizontal gene transfer, providing robust detection sensitivity without excessive false positives.

The analysis applied epidemiological filtering (Hamming distance threshold ≤15) and mutation density analysis (3.0% threshold) to identify potential recombination events across 3,253 EFSA-filtered cgMLST loci. Quality filtering retained samples with *≥*98% cgMLST completeness, ensuring high-confidence epidemiological comparisons suitable for outbreak investigation contexts.

Analysis revealed recombination events in 20 sample pairs, affecting 0.03% of EFSA loci. Notably, all samples involved in recombination events with available metadata were patient samples, while most lacked temporal information, contrasting with 50% of the dataset that includes collection years spanning 1987-2021. Mutation density analysis revealed recombination events with densities ranging from the threshold minimum (3.0%) to high levels, indicating substantial nucleotide-level differences that would be undetectable using traditional cgMLST Hamming distance analysis.

#### Recombination Impact on Clustering Robustness

To assess whether recombination events compromise the reliability of our clustering and source prioritization results, we evaluated the overlap between recombination events and cluster membership. This analysis addresses whether our multi-scale clustering conclusions remain valid when potential recombination events are considered.

In Se-1540, recombination events occurred primarily between samples that were already separated in our clustering analysis, indicating that distance-based clustering naturally segregates samples with extensive horizontal gene transfer from epidemiologically relevant transmission clusters. This demonstrates that our Tier 1-2 analytical framework successfully focuses on epidemiologically relevant genetic variation while maintaining robustness against population-level genetic phenomena.

The minimal overlap between recombination events and established clusters confirms that nucleotide-level analysis provides enhanced discrimination without being compromised by recombination artifacts. This validation supports the reliability of our clustering and source prioritization results for epidemiological outbreak investigation purposes.

The recombination analysis capability represents a preliminary implementation demonstrating cgDist’s extensibility beyond distance calculation to encompass broader epidemiological genomics applications. While this analysis provides proof-of-concept validation of the multi-scale surveillance workflow, comprehensive evaluation of recombination detection accuracy and epidemiological utility would require additional validation studies with experimentally verified recombination events and detailed comparison to established recombination detection methods.

